# Evolution of the skull in arvicoline cricetids (Rodentia) according to 3D morphometric insights: Part 1. Morphological disparity of the palato-spheno-pterygoid complex

**DOI:** 10.1101/2024.09.04.611334

**Authors:** Leonid L. Voyta, Daniel A. Melnikov

## Abstract

Our paper is the first contribution to the comprehensive analysis of the complicated evolution of cranial and mandibular parts connected by pterygoid muscles, as part of more global investigation into adaptive evolution of Arvicolinae. The analysis was performed on 90 micro-computed-tomography–scanned specimens from 33 species from 19 genera of Arvicolinae as well as two species from two genera of Cricetinae as an outgroup. We revised 11 morphological traits of the “palato-spheno-pterygoid” complex, including key features of the palatine that are highly important for defining Arvicolinae taxa according to the micro-computed-tomography data. We also homologized characters of the posterior palatal margin and categorized the composition of the palatal elements into two main morphotypes: morphotype “A” is unique to Clethrionomyini and morphotype “B” was subdivided into three additional types and was found to occur in the outgroup (cricetines), voles (B2), and lemmings (B3). Morphospace analysis of the palato-spheno-pterygoid complex by means of the three-dimensional dataset revealed a mode of transformation of morphotype “A” into morphotype “B2.” A separate task was the development of a protocol for the preparation of morphological data for subsequent evaluation of genotype–phenotype relationships using specialized software applications (e.g., RERconverge).

## Introduction

It is widely recognized that fossils play a crucial role in phylogenetic reconstructions and estimation of species divergence times (Withnell and Scarpetta 2024). For some taxonomic groups, the fossil record is the only way to study them (Michaux et al. 2012; Fabre et al. 2014; Krause et al. 2014; Wang et al. 2019; Mao et al. 2024; and others). Moreover, "in the context of skeleton-based phylogenies, fossils will have a more significant effect if the tree contains long terminal branches and short internal branches" (Smith 1994: 68).

Rapid development of modern genomic and bioinformatic technologies is generating new methods for combining molecular and morphological data (Hiller et al. 2012; Adams 2014; Prudent et al. 2016; Hu et al. 2019; Kowalczyk et al. 2019; Redlich et al. 2023). It is therefore crucial to find ways to describe the morphology of fossil taxa in detail while taking into account physical fragmentation and poor preservation. Furthermore, the morphological data should be formalized in a specific way (e.g., see Kowalczyk et al. 2019; Redlich et al. 2023). A solution to this research problem will require model taxonomic groups with a sufficiently well-developed molecular phylogeny and broad representation in fossil and recent faunas. Arvicolinae (Cricetidae, Rodentia) is such a model group in this context.

Arvicoline rodents represent one of the most outstanding and well-studied taxonomic group of eutherian mammals. There are over 150 extant and even more extinct species that have emerged since the Neogene in the Northern Hemisphere (Musser and Carleton 2005; Fejfar et al. 2011; Pardiñas et al. 2017). In response to global environmental changes in Eurasia and North America, several cricetid lineages evolved during the Neogene as “microtoid cricetids,” which gave rise to true arvicoline cricetids (Fejfar et al. 2011).

The main evolutionary trends traditionally ascribed to voles include an increase in the height of cheek tooth crowns and the development of prismatic molars (Koenigswald 1993; Maul 2001; Fejfar et al. 2011). Nonetheless, there is a notable gap in research on evolutionary changes in cranial and mandibular structures in relation to the evolution of dentition in the taxonomic group. Although classic studies by Repenning (1968), Kesner (1980), and Vorontsov (1982) have provided valuable insights, there is still a need for further research in this area.

The phylogeny of Arvicolinae has been studied by both morphological (Robovský et al. 2008) and genetic (Abramson et al. 2009, 2021; Bannikova et al. 2010) methods, with differing taxonomic results in some cases (Pardiñas et al. 2017; Kryštufek and Shenbrot 2022). The application of molecular phylogenetic methods has generally supported the hypothesis of three successive waves of adaptive radiation (Figure 1) in evolutionary history of the taxonomic group (Abramson et al. 2009, 2021). Of particular interest within the “three-wave radiation” paradigm is clear definition of the Clethrionomyini clade—the second successive wave—which exhibits distinctive phenotypic characteristics related to the shape of the palatal margin (Zazhigin 1980; Pozdnyakov 2008). Nevertheless, possible recapitulation of phenotypic traits within Arvicolinae cannot be detected and explained by classic methods. It seems that the description of specific characteristics (Pozdnyakov 2008; Abramson et al. 2020: 9), rather than functional links between osteological and myological complexes (Repenning 1968; Kesner 1980; Vorontsov 1982), is the main methodological limitation restricting our understanding of the evolving phenotypic traits of the taxonomic group against the background of a well-developed molecular phylogeny.

**Figure 1.**
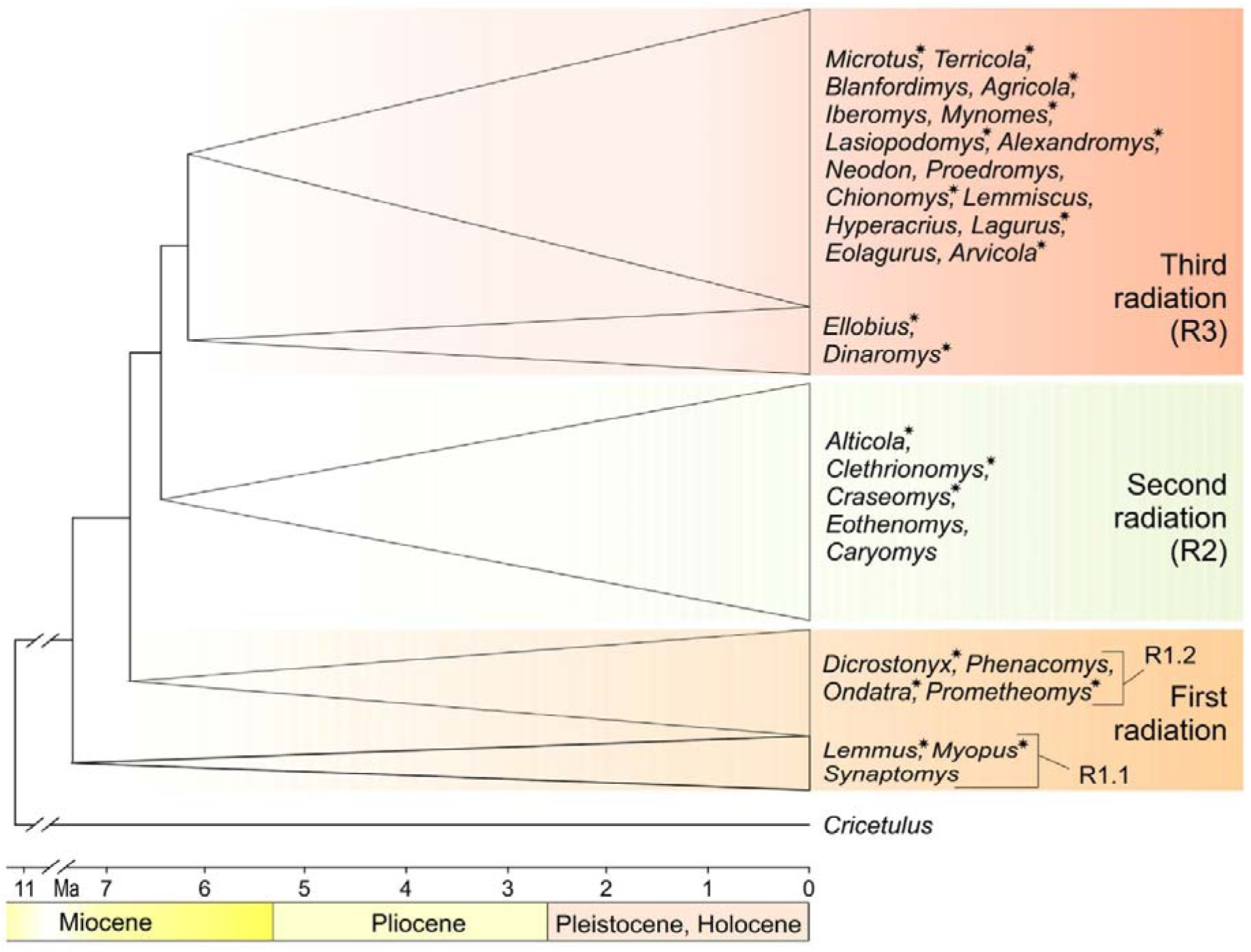
Schematic phylogeny of Arvicolinae based on the time-calibrated mitochondrial genome phylogeny of Abramson et al. (2021). Different colors indicate three successive waves of adaptive radiation in the evolutionary history of the subfamily; star indicates analyzed groups.

Because classic comparative studies on arvicoline palatine traits (e.g., Zazhigin 1980; Pozdnyakov 2008) do not address structural complexity and interbone relations, the aim of the present work was to develop a protocol for analyzing structural variety of the cranial morphological complex, including the palatine bone, in relation to evolutionary trends of three successive waves of vole adaptive radiation as a whole.

This morphological complex of arvicolines was determined here by assessment of a published correlation between pterygoid muscle attachment and hypsodonty stages, as outlined by Repenning (1968) and Kesner (1980). To gain a comprehensive understanding of the evolving traits linking bones, muscles, and hypsodonty level, we conducted detailed examination of the palato-spheno-pterygoid complex (PSP complex; acronyms see at the Appendices bottom).

A fundamental approach to achieving the aim is to carry out a morphospace analysis that involves three-dimensional (3D) data and geometric morphometric methods (Fabre et al. 2014, 2018; Voyta et al. 2022a,b). Morphospace analysis enables the identification of key patterns of morphological disparity across taxa, thereby facilitating a comparison within a morphogenetic (Drake 2011; Renaud et al. 2012; Singh et al. 2017) or phylogenetic (Terray et al. 2022; Voyta et al. 2024) context.

### Morphological background

A researcher’s opinion about the usefulness of the comparative morphological perspective in systematic or evolutionary studies must be based on his/her attitude to answering these fundamental questions: what is the relation between separately described trait complexes, and how should the analysis of these complexes be optimized in modern studies?

The first attempts to answer the first question were made in the middle of the last century. To date, articles by Repenning (1968) and Kesner (1980) remain the only known sources of detailed information on the relation between masticatory muscles and cranial and mandibular characteristics within Arvicolinae.

In the context of the current paper, the focus is on “Pterygoid-Mylohyoid-Hyoid Constrictor Groups” proposed by Repenning, who has analyzed these muscles, which seem to be important for the structure and shape of the mandible. In this group of muscles, that author paid special attention to describing external and internal pterygoid muscles and the digastric muscle (Repenning 1968: 38). Regarding our interest in the palatine shape of the vole, pterygoid muscles may have a direct influence on the modeling of the respective cranial and mandibular regions through so-called “muscle-induced bone plasticity” (Young and Badyaev 2010: 443).

According to anatomical data from Repenning (1968: 38) and Kesner (1980: 213, 214), the external pterygoid muscle connects the alisphenoid bone (muscle origin) and the middle side of the mandibular condylar process (muscle insertion). The internal pterygoid muscle connects the pterygoid (syn. parapterygoid) fossa—composed of the palatine, basisphenoid, and pterygoid bones (origin)—and the ventral margin of the angular process of the mandible (insertion). Therefore, the two pterygoid muscles—*M. pterygoideus externus* and *M. p. internus*—influence the shape of a large cranial region consisting of the palatine, ali- and basisphenoid (and seemingly presphenoid), and pterygoid bones. On the other hand, the muscles also influence the shape of the mandibular ramus through the insertional part. In this regard, Repenning (1968: 43) noted differences in the position of the origin of the external pterygoid relative to the level of the zygomatic arch between mesodontic *Neotoma* (Cricetinae) and hypselodontic *Microtus* (Arvicolinae) owing to increased molar hypsodonty.

Repenning’s extensive comparative material has allowed that author to identify some variations of mandibular morphology among the Arvicolinae. In particular, he has noticed an exception in the shape of the mandibular ramus in *Clethrionomys rutilus* and *C. gapperi* (Clethrionomyini), which does not occur in *C. glareolus*, in which the ascending ramus is typically arvicoline. A similar condition is probably found in lemmings *Synaptomys* and *Myopus* (Repenning 1968: 51). Kesner (1980) also found some variation of the internal pterygoid muscle and its origins in Clethrionomyini, Lemmini, and Phenacomyini. He wrote: “In *Lemmus*, *Clethrionomys* and *Phenacomys* the internal pterygoids resemble the cricetine condition; there is a reduction in the extent of the various aponeuroses, in particular there is a loss of the bipinnate origin of the lateral part. The cricetine-like internal pterygoid reflects the shallower lateral pterygoid fossae found in these genera” (Kesner 1980: 214).

Despite this important contribution, these are only preliminary data that indicate a relation between the tooth developmental pattern (increasing hypsodonty), specific masticatory muscles, and their expression in the bone structure, which, however, do not allow us to generalize their spatial and structural interaction. Nevertheless, previous authors have helped to identify key taxa of voles *s. lato* that deserve close attention. Curiously, one of these taxa has turned out to be Clethrionomyini, which became our model taxonomic group due to its monophyly and distinct position within Arvicolinae clades. Another taxon that deserves attention is the Lemmini tribe.

Modern methods of comparative morphology, extending into analytical approaches of morphometry, allow studies to be optimized in terms of generalizing spatial and structural interactions between trait complexes. For example, Brassard et al. (2020a,b) investigated the architecture of masticatory muscles in domestic dogs via 3D geometric morphometric approaches to show that “muscle size, and thus the attachment area requirements for individual muscles, are likely drivers of mandibular shape” (Brassard et al. 2020b: 308). Similar combined studies have been performed on murid rodents (see Ginot et al. 2018a,b) but not on cricetids, particularly with a taxon set as large as Repenning’s, who analyzed 23 taxa of Cricetinae and 49 taxa of Arvicolinae (Repenning 1968).

The main idea behind the current paper is that the broad generalizations of Repenning and Kesner, together with the methods of Brassard and Ginot, will enable us to obtain new data on the evolution of certain morphological complexes of arvicolines. We do this here via morphospace analysis (see above).

### Research agenda

The agenda of our work can best be described by quoting from the work of Kowalczyk et al. (2019): "A major motivation in evolutionary biology is to determine which genetic changes underlie phenotypic adaptations." In this context, we typically assume that the phenotypic traits undergoing adaptation are on the surface, while the genetic underpinnings are hidden and require close attention. For many mammalian taxa, however, detailed morphological data have not been formalized for the assessment of genotype–phenotype relationships, as implemented in modern techniques such as Coevol (Lartillot and Poujol 2011), Forward Genomics (Hiller et al. 2012; Prudent et al. 2016), PhyloAcc (Hu et al. 2019), and RERconverge (Kowalczyk et al. 2019; Redlich et al. 2023). Thus, a paradoxical conclusion is that the morphology of many well-studied taxonomic groups has not been a subject of detailed description.

For the characterization of the large taxonomic groups, we need to use only general traits such as body mass, brain size, life history, sexual selection, social organization, diet, activity budget, ranging patterns, and climatic variables (Kamilar and Cooper 2013; Borges et al. 2019; Díez del Molino et al. 2023). Nevertheless, some approaches, namely RERconverge (Kowalczyk et al. 2019), offer the possibility of using both categorical and continuous datasets. For example, the conceptual study by the Zoonomia project (https://zoonomiaproject.org) analyzed the association between a genotype and phenotypic traits by means of RERconverge for binary (hibernation, vocal learning) and continuous (brain size) data from 240 mammalian species (Christmas et al. 2023).

Given that Arvicolinae has a well-developed molecular and mitogenomic phylogeny (Abramson et al. 2009, 2021; Bannikova et al. 2010), the agenda of the current study needs to involve a two-step analysis of phenotypic traits to ensure that the obtained results can be subsequently used as continuous data (trait vectors) for searching for a genotype–phenotype relationship using, for example, R package RERconverge (Kowalczyk et al. 2019; Christmas et al. 2023).

The first step of the analysis must be based on investigation of the cranial trait complex, namely the PSP complex, by morphospace analysis. The second step, according to the idea of Repenning (1968), must involve examination of the mandibular ramus in relation to cranial characteristics and myological data. Both steps are carried out in parts 1 and 2 of the present study.

Because morphospace analyses rely on principal component analysis (PCA), each loaded principal component of shape/size variety can be then used as a trait vector via the RERconverge approach or for assessing a phylogenetic signal via the Adams (2014) approach (Terray et al. 2022; Voet et al. 2022). In addition, new information can be obtained by analytically combining the cranial and mandibular trait vectors by a regression analysis (e.g., see Ginot et al. 2018b; Voyta et al. 2024).

## Material and methods

### Sampling

The analysis was performed on 90 micro–computed tomography (CT)-scanned specimens from 33 species from 19 genera of Arvicolinae as well as two species from two genera of Cricetinae as an outgroup (Supplementary Material, Table S1): *Agricola agrestis* (Linnaeus, 1761), *Alexandromys fortis* (Büchner, 1889), *A. middendorffii* (Poljakov, 1881), *A. oeconomus* (Pallas, 1776), *Alticola argentatus* (Severtzov, 1879), *A. lemminus* Miller, 1898, *A. macrotis* (Radde, 1862), *A. semicanus* Allen, 1924, *Arvicola amphibius* (Linnaeus, 1758), *Chionomys nivalis* (Martins, 1842), *Clethrionomys centralis* Miller, 1906, *C. glareolus* (Schreber, 1780), *C. rutilus* (Pallas, 1779), *Craseomys rufocanus* (Sundevall, 1846), *Dicrostonyx torquatus* (Pallas, 1778), *Dinaromys bogdanovi* (Martino, 1922), *Ellobius lutescens* Thomas, 1897, *E. talpinus* (Pallas, 1770), *Lagurus lagurus* (Pallas, 1773), *Lasiopodomys brandtii* (Radde, 1861), *L. gregalis* (Pallas, 1779), *L. mandarinus* (Milne-Edwards, 1871), *L. raddei* (Poljakov, 1881), *Lemmus sibiricus* (Kerr, 1792), *Microtus arvalis* (Pallas, 1778), *Mynomes miurus* (Osgood, 1901), *M. ochrogaster* (Wagner, 1842), *M. pennsylvanicus* (Ord, 1815), *M. richardsoni* (DeKay, 1842), *Myopus schisticolor* (Lilljeborg, 1844), *Ondatra zibethicus* (Linnaeus, 1766), *Prometheomys schaposchnikowi* Satunin, 1901, *Terricola subterraneus* (de Selys-Longchamps, 1836), *Neotoma mexicana* Baird, 1855, and *Cricetulus barabensis* (Pallas, 1773). The material comes from 73 geographical localities. Three specimens—ZIN 34887, 45399, and 83753(6951)—have incomplete label information (Table S1).

The species list was compiled according to the dataset of Abramson et al. (2021). For the morphological comparison, we added *Neotoma* according to the datasets of Repenning (1968) and Kesner (1980). *A. middendorffii* was added to the dataset owing to the quality of the micro-CT data.

### Individual age determination

In this paper, we primarily analyzed mature specimens. In some cases, we extended a group of specimens to include immature ones. The main purpose of age determination (and age-related specimen selection) is related to obtaining a definitive character state and bone shape for correct intergroup comparison. Detailed information on sex and age of each specimen is provided in Supplementary Material (Table S1). Sex was taken from a label. Age groups were defined specifically for this study and are presented in the graphical part of Supplementary Material (Figures S1–S35).

For correct determination of relative age stages, we separated the ingroup dataset into two subsets according to hypsodonty expression: (1) hypselodont species from 13 genera: *Agricola* (Figure S1), *Alexandromys* (Figures S2–S4), *Alticola* (Figures S5–S8), *Arvicola* (Figure S9), *Chionomys* (Figure S10), *Dicrostonyx* (Figure S12), *Lagurus* (Figure S16), *Lasiopodomys* (Figures S17–S20), *Lemmus* (Figure S21), *Microtus* (Figure S22), *Mynomes* (Figures S23–S26), *Myopus* (Figure S30), and *Terricola* (Figure S33); (2) hypsodont species from six genera: *Craseomys* (Figure S11), *Dinaromys* (Figure S13), *Ellobius* (Figures S14–S15), *Clethrionomys* (Figures S27–S29), *Ondatra* (Figure S31), and *Prometheomys* (Figure S32). *Neotoma* bears mesodont teeth (Figure S34), and *Cricetulus* bears brachyodont teeth (Figure S34).

The graphical part of Supplementary Material is Figures S1–S35, showing the skull of each specimen analyzed in two aspects: lateral and dorsal, with age-related features marked and a “local age scale.” The scale is a colored panel at the bottom indicating a relative position of each specimen between five relative age groups: juvenile (juv), immature (sad), mature “ad1,” mature “ad2,” and senile (sen). Juvenile individuals are absent from the dataset. Figures for hypsodont species include radiographs of the root condition of molars. Appendix 1 exemplifies the cranial and dental features used to determine age.

### Species identification and sample sizes

Because the global idea behind the study was to make morphological data suitable for the assessment of genotype–phenotype relationships, it seemed be sufficient to obtain formalized information for each examined species, i.e., each species had to be represented by a single value of a respective trait vector (principal component) obtained from the morphospace. Due to the high taxonomic diversity of the analyzed dataset and the corresponding great morphological disparity within Arvicolinae (19 genera and 33 species of the ingroup), we employed a relatively small sample size for each species, two to six specimens, more often three to four individuals. Seven species were represented a single specimen: *A. middendorffii*, *A. tuvinicus*, *C. rufocanus*, *M. miurus*, *M. ochrogaster*, *M. richardsoni*, and *C. glareolus*. The use of animals of different ages from different geographical localities (for wide-ranging species, see Supplementary Material, Table S1) allowed us to obtain fairly reliable results on the position of each species within morphospace.

Species were determined according to an identification guide (Gromov and Erbajeva 1995). Some identifications were corrected either by experts (Natalia I. Abramson, Semyon Yu. Bodrov, and Feodor N. Golenishchev) in a respective taxonomic group or by means of published studies (Petrova et al. 2022).

### Computed X-ray micro-tomography and segmentation

NeoScan N80 (FP) scanners were utilized to scan the analyzed skulls at the core facility Taxon (http://www.ckp-rf.ru/ckp/3038/) of the Zoological Institute of the Russian Academy of Sciences (St. Petersburg, Russia). Technical specifications are listed in Supplementary Material (Table S2). Specimens were scanned in three-slice mode. Preprocessing was performed in DataViewer software ver. 1.5.4.0 64-bit (SkyScan, Brucker microCT). The main processing of the skulls and of the specific PSP complex was performed using Avizo 2019.1 (FEI SAS). The 3D models, together with additional information, were deposited on the website of research project No. 19-74-20110 (https://zin.ru/labs/evolgenome/morphology/index_Part1.html).

### Morphometric analysis, statistics, and visualization of results

PSP complex shapes were captured as sets of 3D coordinates composed of 71 landmarks (Lms, Figure 2). The complex was described by means of three lines of semilandmarks on the left side: (1) Lms 1–20 along the midline of the dorsal surface of the basisphenoid between the presphenoid–basisphenoid contact (Lm1) and the basisphenoid–basioccipital contact (Lm2); (2) Lms 21–40 along the dorsomedial margin of the pterygoid hamulus (Lm21 at the presumed pterygoid–basisphenoid contact) and the longitudinal lamina of the palatine (the last landmark [Lm21] in location relative to the lng/pal-pm morphotype “A”, see the Results section); (3) Lms 41–70 along the parapterygoid fossa margin from the hamulus (Lm41) to the sphenosquamosal suture (Lm70); and a single true landmark in the posterior nasal spine area (Lm71). Landmarks and semilandmarks were created in the 3D Slicer software ver. 5.4.0 r31938/311cb26 (Fedorov et al. 2012).

**Figure 2.**
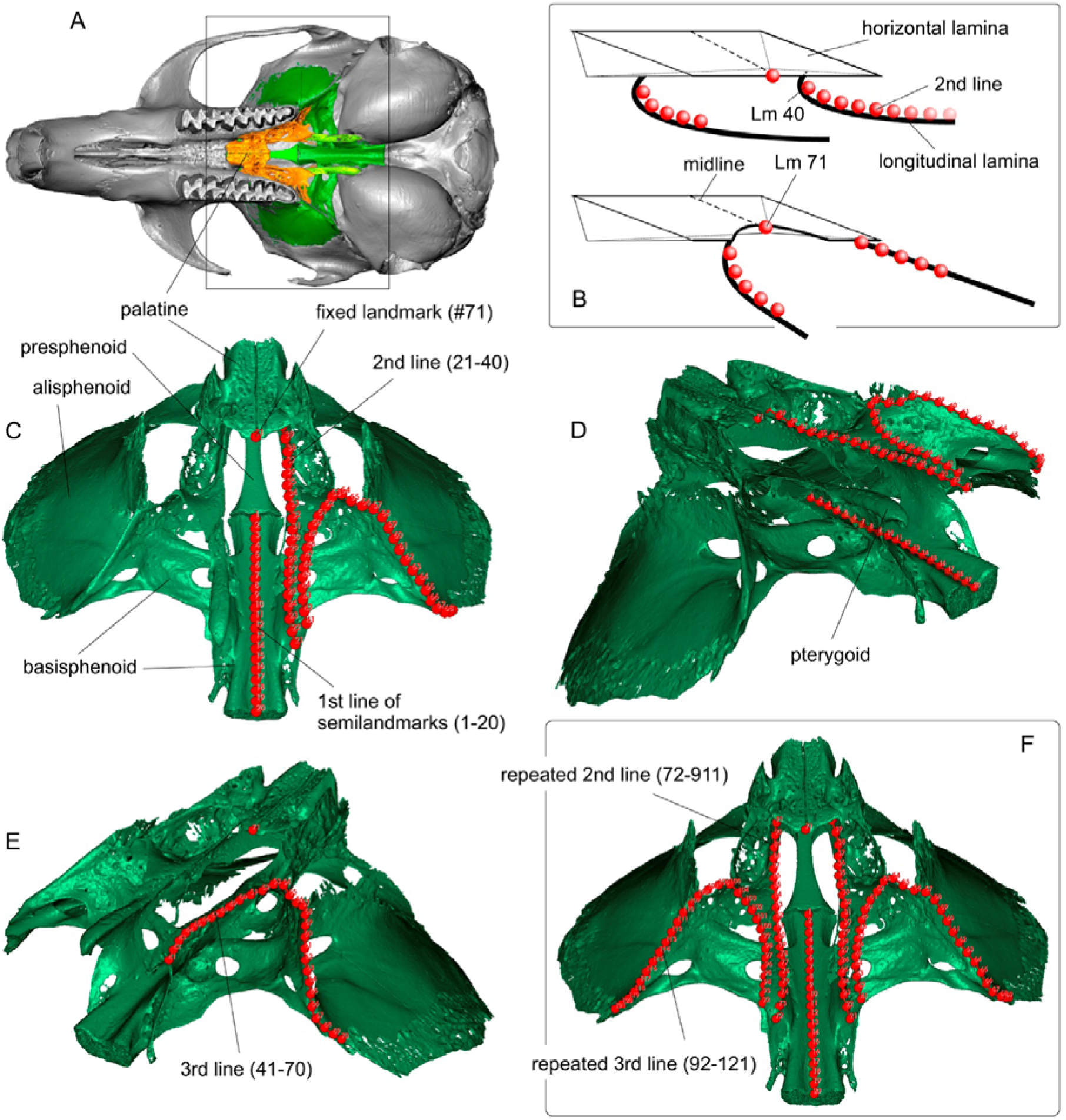
Three-dimensional landmarks position on the left side of the PSP complex (C–E) and expanded landmark wireframe on the right side for visualization purposes (F). (A) Skull of *Alticola argentatus* (ZIN 77489) in ventral view with colored PSP complex (palatine is orange); (B) diagrammatic representation of two main types of the horizontal and longitudinal laminae combining (clethrionomyins type above, arvicolins type below; see details below); (C) screen image of the isolated 3D model of the PSP complex in ventral posterior view (n lms = 71); (D) *ibid*., in ventral posterolateral view; *ibid*., in ventral posterolateral view opposite D image (only the third line is shown; the others were turned off); (E) expanded landmarks set from left to right side (n lms = 121). A made on Avizo, C–E made on 3D Slicer. Not to scale.

Stability and repeatability of landmark positions and semilandmark curves in different groups of specimens were ensured in two ways: (a) continuous comparison of point positions with a reference opened in an adjacent screen of the software (two screens of 3D Slicer); (b) a “measurement error” reduction via landmarking of each specimen group three times, with final PCA of the mean values (see Voyta et al. 2024). Additionally, we tested a variation of the key features of the PSP complex that are associated with the landmark wireframe using image series for each analyzed specimen group (see Supplementary Material, Figures S36–S63). This technique allowed us to assess geographical or individual age differences between specimens of the same species and to avoid overly large discrepancies in landmark positions. Appendix 1 (entry A1) is an example of a wireframe reference for the landmarks of *C. rufocanus*.

The morphometric analysis was performed in three steps. The first step was the Procrustes superimposition procedure (Rohlf 1993; Zelditch et al. 2004), which was performed with the "SlicerMorph" module of the 3D Slicer software (Fedorov et al. 2012), resulting in a Procrustes coordinate matrix. The second step was PCA and "live view" visualization of shape transformation by means of SlicerMorph capabilities. The third step was final PCA analysis and obtaining of morphospace plots using the Procrustes coordinate matrix from the first step and PAST software ver. 4.03 (Hammer et al. 2001). This step ensured the combination of three subsets (landmarking repetitions A–C) for implementing the protocol of the “measurement error” reduction.

Given that the main landmark dataset involved only the left side of the PSP complex, we had difficulties with the final visualization of the shape transformation in morphospace owing to the one-sided distortion of the bilateral symmetric reference model. The decision to use only half of the complex was driven by the preservation of the material because most of a skull was partially damaged, especially pterygoids. To solve this problem, we defined pairs of comparisons using four first principal components (see below for details). Each pair consisted of species from negative and positive ends of a respective principal component. After the pairs were defined, the PCA was repeated on the basis of the reduced dataset using SlicerMorph. The correctness of this approach is supported by theoretical papers (e.g., Rohlf and Slice 1990) and pair comparison protocols realized for example in the Morpho R package (Schlager 2017) [see also ref. (Claude 2008); regarding practice, see refs. (Voyta et al. 2022a,b, 2024)]. Furthermore, we tested the correctness by paired tests performed on two blocks of Procrustes distance values of the full datasets vs. the reduced subset.

The extension of the PSP complex size from unilateral to bilateral required addition of landmarks. The subset was described by means of 71 initial landmarks and 50 additional semilandmarks: repeated second (20 Lms along of the longitudinal lamina) and third (30 Lms) lines (Figure 2F).

Another aspect of the visualization of results is related to 2D and 3D PCA plots. In this study, standard 2D plots were employed with PAST software. To facilitate determination of interspecies differences within complex morphospace, 3D plots were generated using R packages Scatterplot3d (Ligges et al. 2023) and RGL (Adler and Murdoch 2023).

The shape analysis of the PSP complex yielded 89 principal components describing 44.263% (PC1) to 0.000048% (PC89) of the variance. In this case, we should address the question of "how many ordination axes should be considered non-trivial and interpretable" (Jackson 1993: 2204). In our previous published studies (Voyta et al. 2022a,b, 2024), we used the "broken-stick model" from the algorithm of Jackson (1993) to select the axes; the PAST software (Hammer et al. 2001) performs this procedure by default. Nonetheless, we were often faced with “in-between” or ambiguous results, as in this case (Appendix 2A). In the current analysis, “the stick” touches a fifth principal component without clear guidelines as to how to determine an axis count in this case. Therefore, we utilized an additional assessment (“scree test”) from Jackson’s review, which revealed graphical differences between observed (our PCA) and artificial (random) datasets (Horn 1965). The protocol was implemented in the PAST software. The random dataset was represented by a matrix whose row and column numbers corresponded to the observed data (90 rows/213 columns). The random data were generated via Microsoft Excel equation “=RAND()” and used in the standard PCA pipeline. The scree test was performed with the percentage variance of the first 15 axes of the two datasets pooled on the joint bivariate plot (Appendix 2B). By analogy to Horn’s simulation, we obtained an intersection between the sample curve and the random curve through the sixth axis; therefore, the significant axes were the first five principal components. On the basis of the tests, we decided to employ only the first four axes to describe morphological disparity. The controversial fifth axis was excluded because its value differed greatly from that of the fourth axis (3.508% vs. 9.378%). The limit value of the fifth axis was well represented by the broken-stick model (Appendix 2A). Our test comparison suggests an interpretation of such controversial cases as follows: take the axes clearly above the “broken stick.”

### The morphological-description strategy and terminology

The morphological part of the study revised osteological features of the arvicoline PSP complex because the nomenclature and homology of some palatine and sphenoidal features are still undeveloped or unstable. Furthermore, there is no detailed description of bones of the complex, including the palatine, in the literature. Consequently, in the current investigation, we selected the following strategy: (a) use the most detailed published descriptions of rodent bones, (b) use the functional aspect of distinctive features of the PSP complex, and (c) use visualization capabilities of 3D models and specialized software, similarly to Wible (2022).

(a) Modern sufficiently detailed original studies on rodent cranial anatomy are presented in relatively few papers, including the detailed description of Sigmodontinae by Weksler (2006) and Missagia and Perini (2018) and of Murinae by Fabre et al. (2018). It should be noted that the PSP complex is only briefly described there. Only Fabre et al. considered palatal foramina and grooves in sufficient detail. We also followed Wahlert (1974) in describing foramina of the alisphenoid canal complex.

On the other hand, in the context of my (L.L.V.) core competence in Soricomorpha, several Wible papers should be mentioned, first of all the description of the solenodon (Wible 2008) and the modern description of the gray short-tailed opossum (Wible 2022) as well as ctenodactyloid rodents (Wible et al. 2005). In the papers, that author refers throughout to a canonical anatomical description of the dog by Evans (1993) and to classic articles by McDowell (1958), Novacek (1986), and MacPhee (1981, 1994), which allowed specimens to be examined in the broader context of intergroup homologies and functional relations between different morphological complexes. Furthermore, the link between Wible’s papers and "Miller’s Anatomy of the Dog" (Evans 1993) indicates universality of the anatomical nomenclature.

Therefore, the anatomical nomenclature and the topology of the morphological characteristics were primarily in accordance with those of Wible’s studies (Wible et al. 2005; Wible 2008) and a guideline by Evans (1993). The aforementioned rodent studies (Weksler 2006; Fabre et al. 2018; Missagia and Perini 2018) were utilized to assess intergroup variation of characteristics.

(b) The functional aspect is related to "topological criteria" in the sense of Van Valen (1994). According to Van Valen (1994: 153), "topology, which should not be confused with topography, comprises those relations of the elements of a space which are preserved under continuous deformations, such as change of size or shape." That author himself used the term primarily for dental homologies. For bone homology, nutritional and innervation tracings are also important. In this regard, we used specialized angiological papers on rodents such as (Bugge 1970) and (Brylsky 1990), with some corrections by Evans (1993). A paper directly about palatal surface structure was published by Quay (1954).

This aspect is also related to myology, which we consider through the aforementioned papers of Repenning (1968) and Kesner (1980) as well as some modern articles (Hautier 2010; Baverstock et al. 2013).

(c) In the current work, we made extensive use of 3D skull models and separate reconstruction (segmentation) of individual bones. Due to differences in the bone preservation state, it was not possible to reconstruct the PSP complex of all species at the same level of detail. Therefore, only three skulls were processed at a high level of detail—*A. argentatus* (ZIN 77489), *A. middendorffii* (ZIN 101018), and the outgroup species *C. barabensis* (ZIN 101697)—to prove "phylogenetic criteria" of homology (MacPhee 1981). Other specimens were described by means of entire 3D models using "surface cross contour” and “surface cross section” functions of Avizo and radiographic data as a byproduct of micro-CT.

## Results

### Palatine

The palatine of Arvicolinae is a relatively long bone featuring maximum contact with the maxilla, moderate contact with the sphenoid, and limited contact with the frontal, ethmoid, and vomer at the rostral end as well as the pterygoid at the caudal end. In general, the palatine is composed of three laminae: the horizontal, perpendicular, and sphenoethmoidal (Evans 1993: 152). The latter is undeveloped in arvicolines because of the narrow connection of the palatine to the ethmoid and vomer. The bone was also found to contain maxillary and sphenoidal processes or facets (Figures 3 and 4).

**Figure 3.**
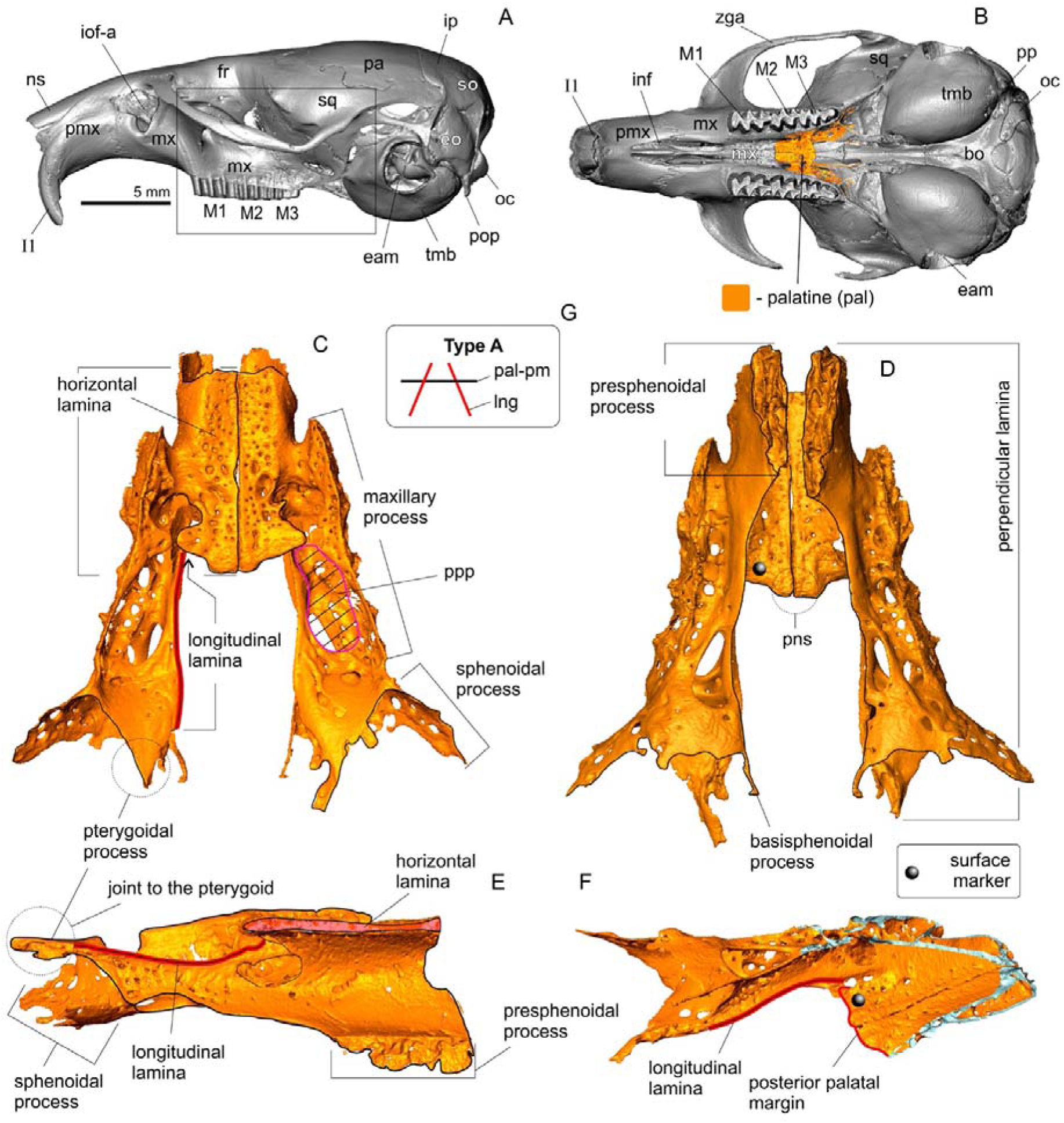
The position of the palatine on the skull of *Alticola argentatus* (ZIN 77489) with marking of anatomical structures and detailed description of the isolated palatine. (A) Skull in lateral view; (B) skull in ventral view; (C) isolated palatine in ventral view; (D) isolated palatine in dorsal view; (E) right palatine bone in medial aspect; (F) cropped palatine part in ventral medial aspect (see ‘surface marker’); (G) pictogram of morphotype A inherent Cletrionomyini. Not to scale. *Key*: **bo**, basioccipital; **bs**, basisphenoid; **eam**, external acoustic meatus; **eo**, exoccipital, **fr**, frontal; **ham**, pterygoid hamulus; **I1**, first upper incisive; **inf**, incisive foramen; **iof-a**, infraorbital foramen, anterior opening; **ip**, interparietal; **lng**, longitudinal lamina (lamellar ridge); **M1–M3**, upper molars (1–3); **mx**, maxilla; **ns**, nasal; **oc**, occipital condyle; **pa**, parietal; **pal**, palatine; **pal-pm**, palatal posterior margin; **pmx**, premaxilla; **pns**, posterior nasal spine; **pp**, paraoccipital process; **ppp**, posterolateral pit; **sq**, squamosal; **tmb**, tympanic bulla; **zga**, zygomatic arch.

**Figure 4.**
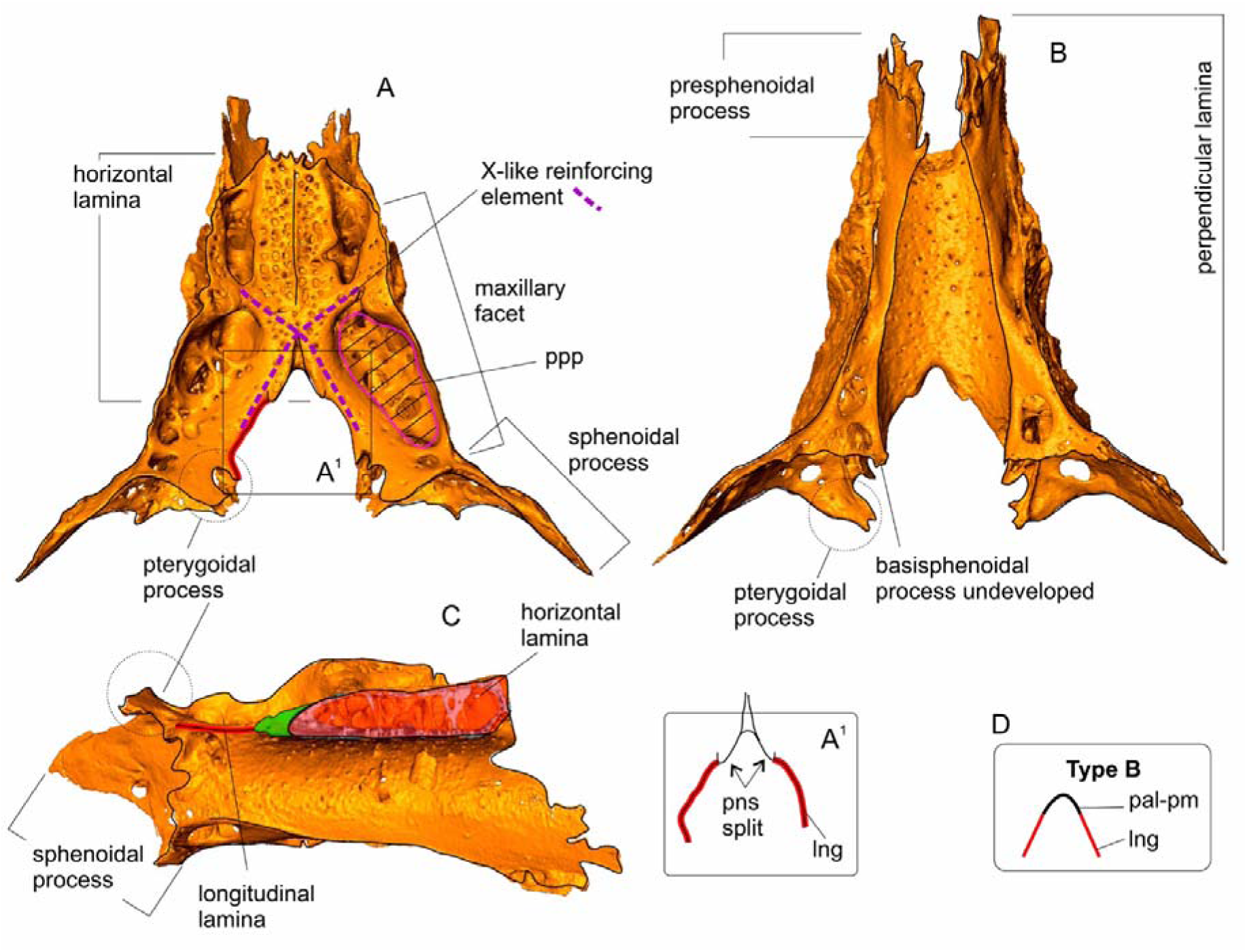
Isolated palatine of *Alexandromys middendorffii* (ZIN 101018). (A) Isolated palatine in ventral view; (A^1^) explanatory drawing of the fused longitudinal lamina and the split posterior nasal spine; (B) isolated palatine in dorsal view; (C) right palatine bone in medial aspect; (D) pictogram of morphotype B. Not to scale. *Key*: **lng**, longitudinal lamina (lamellar ridge); **pal-pm**, palatal posterior margin; **pns**, posterior nasal spine; **ppp**, posterolateral pit.

The palatine of arvicolines, just as that of cricetines, differs significantly from the bones of the model mammals, e.g., dog (Evans 1993: 152) or solenodon (Wible 2008: 335), in that the caudal part of the perpendicular lamina has strongly developed into a huge parapterygoid fossa. The palatine of *C. barabensis* has the shallowest parapterygoid fossa among the analyzed species, but its area exceeds the area of the horizontal lamina (Appendix 3). *Neotoma* has an elongated and relatively deep fossa (Figure S63).

The evolution of hypsodonty and complexity of chewing movements in voles *s. lato* significantly modified the caudal part of the perpendicular lamina of all taxonomic group members. In comparison to hamsters, these modifications have extended the lamina of voles in two directions: (i) the angle of the parapterygoid fossa changed from subhorizontal in hamsters to almost vertical in voles. The functional aspect of this change was discussed by Kesner (1980: 214), who noted that "shifts in orientation of the attachments of the internal pterygoid [muscle] in microtines bring the muscle more nearly parallel with the frontal plane through the molar tooth row than in cricetines, thus reducing the vertical vector component and increasing the anterior and medial components of this muscle"; (ii) there is a longitudinal extension of the intermediate part of the perpendicular lamina between the level of the posterior palatal margin and the anteroventral margin of the parapterygoid fossa (Appendix 4C) in relation to the increase in the length of the arvicoline tooth row and in particular the enlargement of the individual teeth. Moreover, the two aforementioned modifications greatly transformed the bone and added new distinctive morphological elements {1} that are currently undescribed in the model animals (i.e., dog, solenodon, opossum, and ctenodactyloids) and {2} whose nomenclature is undeveloped.

The first additional characteristic is the pterygoidal process of the perpendicular lamina (Figure 3). The process is well developed in Clethrionomyini (Figures 3, S11, S8, S39–S42, S57, and S58), *Prometheomys* (Figure S61), and *Cricetulus* (Appendix 3) and poorly developed in *Lemmus* (Figure S54), *Myopus* (Figure S59), and *Neotoma* (Figure S63). Other examined species showed moderate development of the pterygoidal process of the palatine (Figure 4).

The next new characteristic is a thin basisphenoidal process, but it was most often found to be broken off. The process contacts the most posteromedial tip of the perpendicular lamina and the base of the pterygoid process of the basisphenoid. This characteristic is described here for *Alticola* (Figure 3) and *Cricetulus* (Appendix 3).

An additional characteristic is a longitudinal lamina that forms a ventral margin of the mesopterygoid fossa and occupies a relatively wide space between the fossa (medially) and the lateral margin of the posterolateral palatal pit, at some distance from the palatomaxillary contact (Figure 3). The caudal end of the lamellar ridge is in contact with the pterygoid (Appendix 3C), but the rostral end varies among different taxonomic groups thus forming two main morphotypes (Appendix 4D). Morphotype “A” here was described here for *Alticola* (Figure 3) and represents a conventionally independent position of the posterior palatal margin and of the lamellar ridge: the ridge lies dorsal and rostral of the palatal margin. This morphotype was found to occur exclusively in Clethrionomyini (Figures 3, S39–S42, S57, and S58).

Morphotype “B,” described here for *Alexandromys* (Figure 4), represents close contact between the posterior palatal margin and the lamellar ridge: the ridge has fused with the palatal margin, thereby forming a horseshoe-like pattern (Appendix 4D). The morphotype is widespread among arvicolines. Nonetheless, several variants were noted. Morphotype “B1” has primitive hamster-like composition of the palatal margin and of the lamellar ridge: *Cricetulus* and *Neotoma* (Appendix 3; Figure S63). This morphotype differs from the advanced arvicoline state by small thickness of the horizontal lamina (Figure 3C cf. Appendix 3C).

Morphotype “B2” is typical of most tribes of arvicolines, except lemmings *s. lato*. Morphotype “B3” possesses the most advanced composition of the above-mentioned elements when high inflation of the horizontal lamina provides a pattern similar to morphotype “A” (the lamellar ridge was above the palatal margin): *Dicrostonyx*, *Lemmus*, and *Myopus* (Figures S45, S54, and S59). This composition has apparently been achieved in lemmings via overgrowth of the material of the "posterior nasal spine" (Wible 2008: 337) and neighboring tissue.

The palatine of arvicolines shows some other characteristics, especially at the posterior margin of the horizontal lamina, such as the presence or absence of the aforementioned posterior nasal spine. At first glance, we should be describing interspecies differences in spine morphology because some species manifested remarkable expression of this element, e.g., the palatine of *Ellobius* (Appendix 5) or *Ondatra* (Figure S60). Nevertheless, this trait’s expression is subject to considerable variation and requires further study.

On the other hand, in the Cricetinae–Arvicolinae lineage and probably between morphotypes A and B, a significant trend of palatal evolution seems to be general inflation of horizontal and longitudinal laminae. At least members of Clethrionomyini as bearers of morphotype “A” are characterized by a noninflated horizontal lamina (Figure 3E), just as *Cricetulus* (Appendix 3C) and *Neotoma* being closer to primitive morphotype “B1.” The inflation of the elements is also accompanied by a strengthening of the entire structure of the laminae owing to the thickening of the ridges. For example, *A. middendorffii* in Figure 4 shows (i) dense fusion of each half of the posterior nasal spine with the lamellar ridge and (ii) fusion of structures of the longitudinal and horizontal laminae resulting in an X-like powerful element (Figure 4A), which is quite a common feature of arvicolines as a whole. In accordance with the Repenning’s (1968) and Kesner’s (1980) results, we associate this transformation with the transition of arvicolines to advanced "propolinal" (anterior–posterior) movements of the mandible during chewing (Kesner 1980: 216). Certainly, the advantage of voles in propolinal movements, together with the shift to graminivory and progressive hypsodonty, determines a whole spectrum of palatal transformations: from the inflation of the palatal laminae and the X-like reinforcing elements to the taxon-specific alteration of the posterior palatal margin.

### Palatine foramina and grooves

A distinctive feature of voles, compared to hamsters for example, is the complexity of the surface of the hard palate. The ventral surface of the horizontal palatal lamina of arvicolines is riddled with perforations, longitudinal grooves, bridges, and holes. It is likely that these visible features are related to the overall increase in thickness of the lamina as hypsodonty has progressed. Nonetheless, given that purely hypselodont lemmings (Figures S54, S59) have a simpler palatal surface than does root-bearing hypsodont *Clethrionomys* (Figure S57), there is no direct relation with hypsodonty alone.

Using a functional criterion, we can point to only three features that should be discussed: (a) large foramina (at the rostral margin of the horizontal lamina) often accompanied by deep grooves directed rostrally, (b) grooves behind the first foramina, and (c) large pits of the longitudinal lamina.

The first, and perhaps only, detailed description and imaging of the palatal foramina and grooves of voles from a functional point of view was published by Quay (1954). In the *Ondatra* model, that author demonstrated several features important for our study: “P.P.F., posterior palatine foramen,” “P.G., palatine groove,” “Palatine canal,” “P.A. & N., palatine artery and nerve,” and “Sphenorbital fissure” (Quay 1954: 34). The most important part of Quay’s article is the description of the topology of blood vessels, such as the internal maxillary artery (IMA) with its branches, the “palatine artery,” and the sphenopalatine artery.

In this section, we used Quay’s results, his schematic representation of the topology of the vessels and foramina, together with data from the model animals (i.e., dog, solenodon, opossum, and ctenodactyloids), and micro-CT data on homology of the above features within the ingroup compared to the outgroup.

(a) The large foramen at the rostral margin of the horizontal lamina in our data is the "major palatine foramen" in the sense of Evans (1993) and Wible (2008), sometimes without a distinctive anterior groove (Appendix 4A) or with a visible "major palatine groove" (Appendix 4B). The exact definition of this foramen is based on the topology of the palatine canal and passage of the major palatine branch of the IMA (Evans 1993: 152; = major palatine artery) from the "pterygopalatine fossa" (Appendices 6 and 7). Therefore, the major palatine foramen is synonymous with Quay’s posterior palatine foramen, the major palatine groove is synonymous with Quay’s palatine groove, and the major palatine branch of the IMA is synonymous with Quay’s palatine artery (Quay 1954: 34).

The topology of the palatine canal and consequently of the major palatine branch of the IMA (MPA^R^ in the images) was reconstructed by means of micro-CT data and capabilities of the Avizo segmentation tool. This reconstruction is given in Appendix 6 for the reference specimen of *A. argentatus* (ZIN 77489). We also supposed and drew the passage of the main branches of the IMA on the basis of a cast of grooves on the internal surface of the pterygopalatine fossa and spatial composition of the foramina forming the “alisphenoid canal complex.” On the basis of dissection of fluid-preserved specimens of *A. middendorffii* and *Phodopus sungorus* (unpublished data), we propose that the IMA enters through the “caudal opening of the alisphenoid canal,” which is visible in the ventral aspect (Appendix 6B). This topology is supported by an article by Brylski (1990: 35) who wrote: “the internal maxillary artery branches from the external carotid [artery] in the vicinity of the lingual and facial arteries, runs medially to the angle of the mandible, and enters the braincase by the alisphenoid canal [caudal opening of the alisphenoid canal].”

The clear-cut groove cast outlines from the rostral opening of the alisphenoid canal to the palatine canal allowed the passage of the major palatine artery through the pterygopalatine fossa to be confidently reconstructed, and the same was true for the groove cast of the infraorbital branch of the IMA directed laterally around the alveolar capsule of M3 (Appendix 6D). These conclusions hold true for the other analyzed species.

The reconstruction of the mandibular branch of the IMA (Appendix 6D) was only tentative because the fluid-preserved specimens did not yield clear data on their topology in relation to the foramina. In addition, the findings of Bugge (1970) indicate the absence of some arteries, such as the mandibular branch of the IMA in lemmings and *Neotoma*, and the development of anastomoses "between the distal end of the external carotid artery and the proximal part of the mandibular branch [of the IMA]" (Bugge 1970: 322).

(b) Unfortunately, Quay did not analyze the minor palatine branch of the IMA, which forms the "minor palatine foramen" in the dog and solenodon (Evans 1993; Wible 2008). Furthermore, we are not aware of any published data on the minor palatine branch of cricetids. A detailed description of the mammalian models suggests that "the minor palatine nerve and artery arise in the pterygopalatine fossa and run anteroventrally in a deep notch in the posterior edge of the palate between the palatine and maxilla that rarely is closed to form a foramen often accompanied by deep grooves directed rostrally" (Wible 2008: 391).

Nevertheless, the orbital space of cricetids differs significantly from that of the model animals (dog and solenodon) because cricetids feature an extremely posterior shift of the upper dentition in relation to the alisphenoid canal and the sphenorbital fissure. Accordingly, in cricetids, especially arvicolines, the shape of the pterygopalatine fossa is significantly altered, as is the topology of key foramina such as the alisphenoid canal and the foramen ovale. The hamster pterygopalatine fossa is a relatively wide space with a visible position of the dorsal entrance of the palatine canal, whereas all arvicolines have a narrow deep fossa, not visible from the orbit owing to huge alveolar capsules of the second and third upper molars (Appendices 6 and 7). Therefore, we can hypothesize a difference in the origin of the minor palatine branch among arvicolines, but we can be fairly confident in the reconstruction of its entry and passage within the palatal space.

According to the anatomy of the model animals, the artery enters the palatal surface through the posterolateral corner of the palate between the palatine and the maxilla. In our material, we notices a structure (“semicanal” of the minor palatine artery in *Cricetulus)*: a “groove/canal of the minor palatine artery” (Appendix 4: A1). This groove is synonymous with Fabre’s “psu, palatine sulci,“ which he imaged in *Halmaheramys* and which is visible in the same image for *Rattus* (Fabre et al. 2018: 196). A similar groove (sulcus *sensu* Fabre) is present in *Neotoma* (Fig. S63; an obvious state in ZIN 39375) but not in arvicolines.

The next groove associated with the passage of the minor palatine branch is a more or less pronounced longitudinal groove behind the major palatine foramen and in front of an entrance of the artery into the palatal space. This groove is interpreted here as a “minor palatine groove” and is illustrated here in *Cricetulus* (Appendix 4: A1) but is also well-pronounced in both specimens of *Prometheomys* (Figure S61). Other arvicolines show a hypertrophic pit (see below), which probably subsumes the groove.

(c) A large pit in the longitudinal lamina of the palatine is the "ppp, posterolateral pit" of Weksler (2006: 36), who described it for oryzomine rodents (Sigmodontinae, Cricetidae). The pit is absent in *Cricetulus* (Appendix 3) and is very poorly developed in *Neotoma* (Figure S63) but is a typical feature of arvicolines as a whole, among which it is well developed. The pit is located on the longitudinal lamina of the palatine and is usually riddled with perforations, pores, and holes of various sizes (Figures 3 and 4). A fairly common state of pit development in arvicolines is as follows: the rostral extrusion forms a “groove-like” pit. In the case of overgrowth of the horizontal lamina surface, a groove-like pit can be covered by a “bony bridge,“ and thus the groove-like pit becomes a “channel-like” pit.

### Pterygoid

The pterygoid was tentatively delimited as a separate element in some immature specimens at the contact with the pterygoid process of the palatine and was only slightly delimited along the ventral contact with the pterygoid process of the basisphenoid. On the basis of a faint trace of the suture, we reconstructed a separate pterygoid for the reference specimen of *A. argentatus* (Figure 5). Nonetheless, the pterygoid was usually densely fused with the basisphenoid and was not well-pronounced as a separate element.

**Figure 5.**
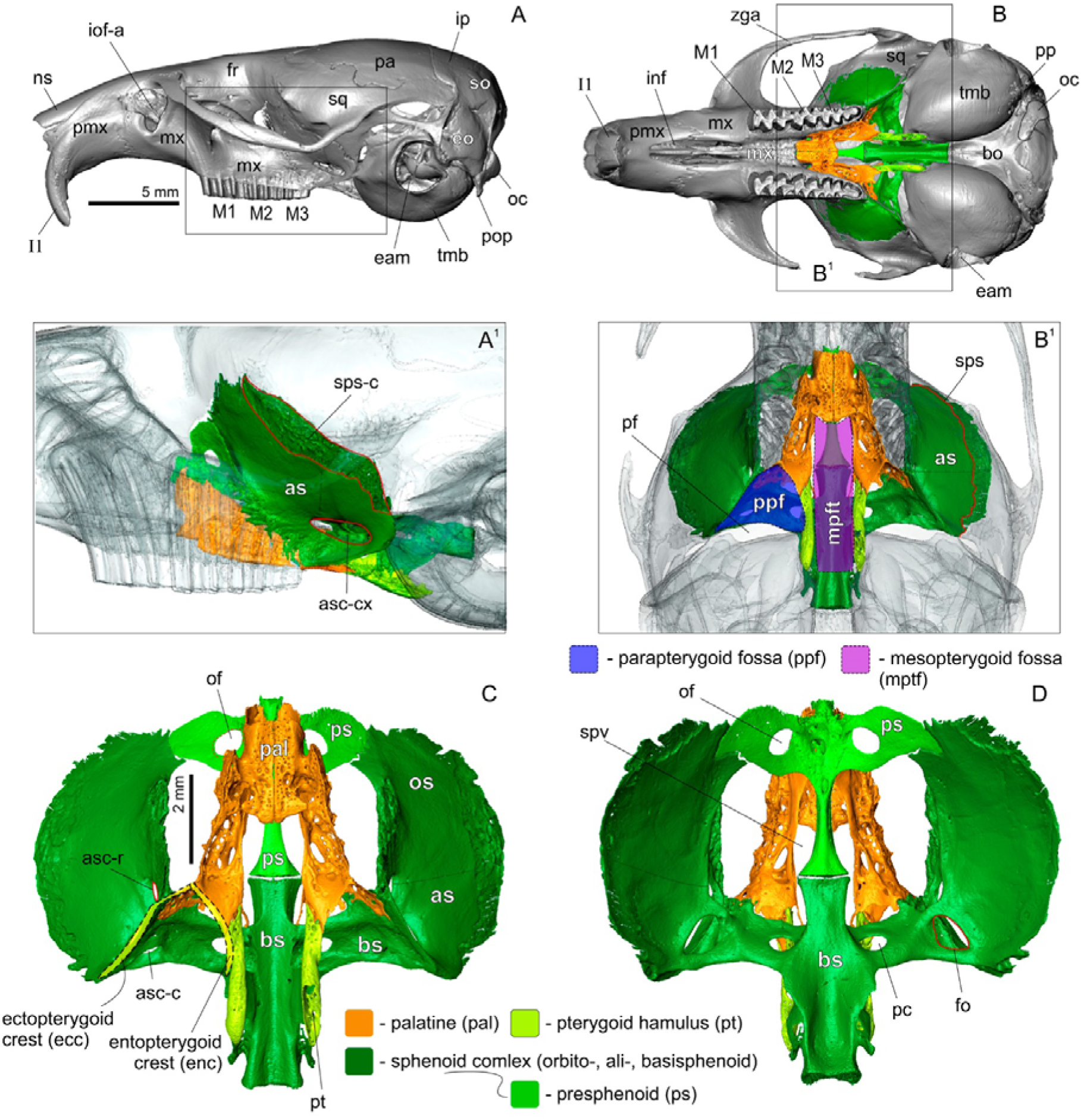
Position of the PSP complex on the skull of *Alticola argentatus* (ZIN 77489) with marking of anatomical structures. (A) Skull in lateral view; (A1) magnified part of the orbital region of the skull (adjacent bones of the PSP complex shown in transparent view: A1 and B1); (B) skull in ventral view; (B1) magnified part of the area of the PSP complex; (C) isolated PSP complex, in ventral view; (D) isolated PSP complex, view on inner surface. A1 and B1 not scaled. *Key*: **asc-c**, caudal opening of the alisphenoid canal; **asc-cx**, alisphenoid canal complex; **asc-r**, rostral opening of the alisphenoid canal; **bo**, basioccipital; **bs**, basisphenoid; **eam**, external acoustic meatus; **ecc**, ectopterygoid crest; **enc**, entopterygoid crest; **eo**, exoccipital, **fr**, frontal; **inf**, incisive foramen; **iof-a**, infraorbital foramen (anterior opening); **ip**, interparietal; **M1–M3**, upper molars (1–3); **mptf**, mesopterygoid fossa (basipharyngeal canal by Wible 2005); **mx**, maxilla; **ns**, nasal; **oc**, occipital condyle; **of**, optic foramen; **os**, orbitosphenoid (fused with alisphenoid); **pa**, parietal; **pal**, palatine; **pc**, pterygoid canal; **pf**, piriform fenestra; **pmx**, premaxilla; **pp**, paraoccipital process; **ppf**, parapterygoid fossa; **ps**, presphenoid; **pt**, pterygoid; **sps**, sphenoparietal suture; **sps-c**, sphenoparietal suture contact area (surface rugosity); **sq**, squamosal; **spv**, sphenopalatine vacuity; **tmb**, tympanic bulla; **zga**, zygomatic arch.

The pterygoid, together with the pterygoid process of the basisphenoid, forms the posterolateral wall of the parapterygoid fossa.

### Sphenoid complex

The complex is composed of four elements, of which the presphenoid and the basisphenoid are more or less clear-cut, at least in the reference specimens of *A. argentatus* (Figure 5), *A. middendorffii* (ZIN 101018), and *C. barabensis* (ZIN 101697). The orbitosphenoid and alisphenoid (sphenoid ala) together form walls of the orbitotemporal region and the floor of the middle cranial fossa (Wible 2008) and are fully fused. Functionally, the basisphenoid is separated from the alisphenoid by the lateral wall of the parapterygoid fossa, but there is no suture.

The presphenoid is the simplest element of the sphenoid complex. The rostral part forms a large orbital wing with a large round optic foramen. The body of the presphenoid is narrow and gradually widens toward distal ends. Anteriorly, the presphenoid is in contact with the ethmoid and vomer, whereas posteriorly, it comes into contact with the basisphenoid.

The basisphenoid in its main plane has a cruciform shape: a longitudinally oriented body and broad lateral processes. The body of the basisphenoid is flat. It touches the presphenoid anteriorly and the basioccipital posteriorly. The lateral process extends to the alisphenoid and contains two large foramina: the medial foramen is defined here as the “pc, posterior opening of the pterygoid canal” (Wible 2008); the lateral foramen is defined here as the caudal opening of the alisphenoid canal (Figure 5D; Appendix 6).

Pterygoid processes of the basisphenoid arise at the base of the lateral processes and are directed ventrally toward the pterygoids. Additionally, at the anterior base of the lateral processes, the basisphenoid body is pierced by a large “trc, transverse sinus canal” (Wible 2008).

The alisphenoid canal complex of arvicolines combines three foramina such as the caudal opening of the alisphenoid canal through which the internal maxillary artery enters the canal, the rostral opening of the alisphenoid canal through which the artery leaves the canal and enters the pterygopalatine fossa; and the foramen ovale, which is located on the internal wall of the canal and transmits the “mandibular branch of the trigeminal nerve and a minute meningeal artery” (Wahlert 1974: 353). The nerve it contains comes from the cranial cavity, while the meningeal branch of the internal maxillary artery is directed into the cavity (Evans 1993). According to this topology, the caudal opening (asc-c) can be found on the lateral process of the basisphenoid in the ventral aspect (Figure 5C; Appendix 6B); the foramen ovale can be seen from inside the cranial cavity in front of the piriform fenestra (Appendix 6D). Both alisphenoid canal openings and the foramen ovale can be seen together in a specially rotated skull (Appendix 6E).

The topology of the alisphenoid canal is very important for the homology of the foramen ovale of arvicolines toward cricetines. Voles and lemmings have a single lumen of the alisphenoid canal, whereas hamsters have two (Appendices 7C and 8; see the Discussion section).

In the current study, the alisphenoid is of interest in relation to the formation of the lateral wall of the parapterygoid fossa and the ectopterygoid crest (in part with the basisphenoid; Figure 5), a location of the facet of the external pterygoid muscle (Appendix 3D), and the alisphenoid canal complex. The first two features certainly influence the shape transformation of the alisphenoid in relation to the evolution of the pterygoid muscles within arvicolines. Thus, the alisphenoid was used in the morphometric analysis.

### Analysis of PSP’s complex shapes

The morphospace analysis of the PSP complex’s shape by means of the set of 71 landmarks revealed two main groups. The first group includes most of arvicolines, except for two tribes, Clethrionomyini and Prometheomyini, which together with the hamster taxa form the second group. Because the second group includes cricetines, we propose to define their shape state as the “ancestral state” (Figure 6).

**Figure 6.**
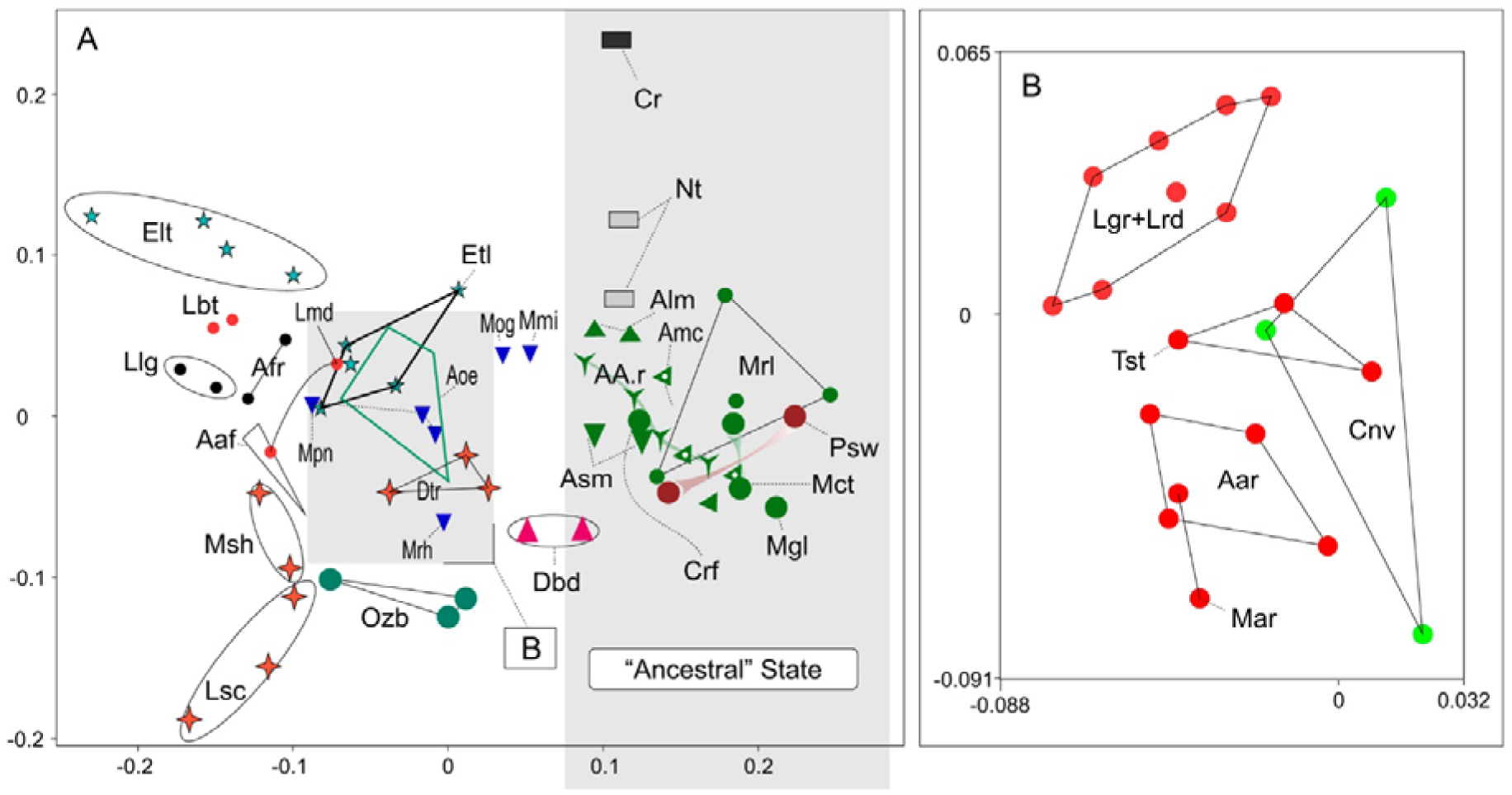
Results of the PSP complex shapes analysis of 33 species from 19 genera of the Arvicolinae and two species from two genera of the Cricetinae (n = 90) based on 3D landmarks data set. (A) Morphospace along of PC1–PC2 axes (axes describe 44.26% of the total variance); (B) enlarged the space area shown the convex-hulls of several species: *Agricola agrestis* (Aar), *Chionomys nivalis* (Cnv), *Lasiopodomys gregalis* (Lgr), *L. raddei* (Lrd), *Microtus arvalis* (Mar), *Terricola subterraneus* (Tst). *Key*: **Aaf**, *Arvicola amphibius*; **AAr**, *Alticola argentatus*; **Afr**, *Alexandromys fortis*; **Alm**, *A. lemminus*; **Amc**, *A. macrotis*; **Amd**, *A. middendorffii*; **Aoe**, *A. oeconomus*; **Asm**, *A. semicanus*; **Crf**, *Craseomys rufocanus*; **Cr**, *Cricetulus barabensis*; **Dbd**, *Dinaromys bogdanovi*; **Dtr**, *Dicrostonyx torquatus*; **Elt**, *Ellobius litescens*; **Etl**, *E. talpinus*; **Lbt**, *Lasiopodomys brandtii*; **Llg**, *Lagurus lagurus*; **Lmd**, *L. mandarinus*; **Lsc**, *Lemmus sibiricus*; **Mct**, *Clethrionomys centralis*; **Mgl**, *C. glareolus*; **Mmi**, *Mynomes miurus*; **Mog**, *M. ochrogaster*; **Mpn**, *M. pennsylvanicus*; **Mrh**, *M. richardsoni*; **Mrl**, *C. rutilus*; **Msh**, *Myopus schisticolor*; **Nt**, *Neotoma mexicana*; **Ozb**, *Ondatra zibethicus*; **Psw**, *Prometheomys schaposchnikowi*; **Tst**, *Terricola subterraneus*.

The positive region of the first principal component (loading was 44.26% of the total variance) occupied the second group with the clethrionomyins core. The negative region of this principal component was occupied by members of tribes Arvicolini, Lagurini, and Ellobiusini and by *Arvicola* together with lemmings and *Ondatra*. The intermediate position was occupied by two species of *Mynomes* (*M. ochrogaster* and *M. miurus*) and by *Dinaromys*.

The second principal component (15.44%) within the first group described differences between *E. lutescens* and *L. sibiricus* specimens, which occupied extreme positive and negative regions of the axis. The second group was segregated into hamsters and clethrionomyins/*Prometheomys* subgroups, so that the variance of PC2 was partly absorbed by the outgroup variation (Figure 6). Figure 6 shows separation of Clethrionomyini with the involvement of *Prometheomys*. To elucidate a relation between these subgroups, we obtained 3D morphospace (PC1–PC3) using R package Scatterplot3d (Ligges et al. 2023), using the subset of PCA values combined into centroids for each species (Table 1). The result is displayed in Figure 7. The figure shows significant separation of cricetines and arvicolines along of the second principal component (Figure 7: B, D) and differences between clethrionomyins and *Prometheomys* along the third principal component (Figure 7C).

**Figure 7.**
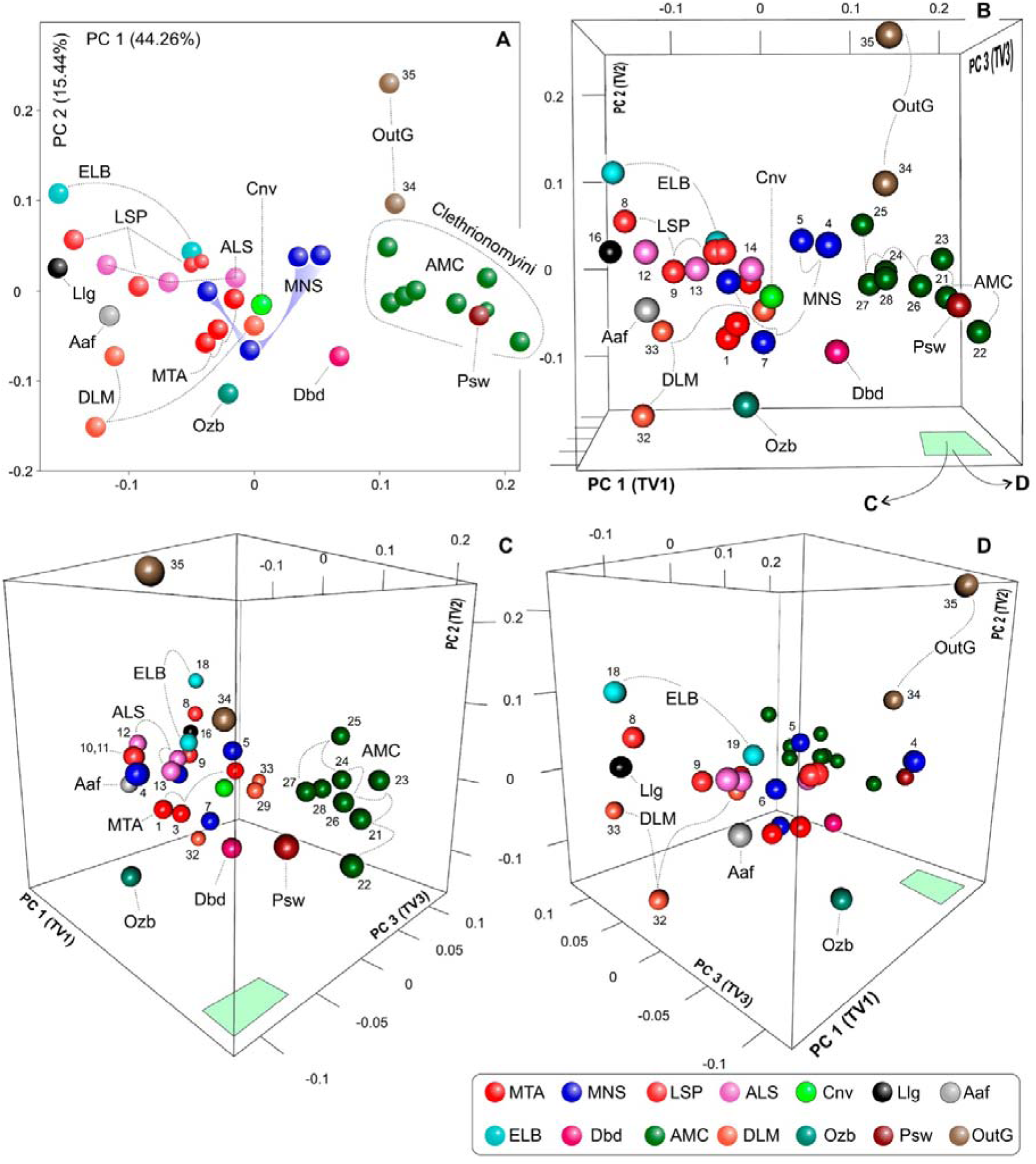
Results of the PSP complex shape analysis shown on the 3D scatter plots (PC1–PC3). The plots obtained on the subset of PCA values (Table 1). (A) 2D morphospace (PC1–PC2); (B) 3D plot in a similar view of image A; (C) 3D plot rotated clockwise (see green panel at bottom); (D) 3D plot rotated counterclockwise. *Key*: **Aaf**, *Arvicola amphibius*; **ALS***, A. fortis* (species ID 12 in Table 1) + *A. middendorffii* (13) + *A. oeconomus* (14); **AMC**, *C. centralis* (21) + *C. glareolus* (22) + *C. rutilus* (23) + *A. argentatus* (24) + *A. lemminus* (25) + *A. macrotis* (26) + *A. semicanus* (27) + *C. rufocanus* (28); **Cnv**, *Chionomys nivalis*; **Dbd**, *Dinaromys bogdanovi*; **DLM**, *D. torquatus* (29) + *L. sibiricus* (32) + *M. schisticolor* (33); **ELB**, *E. lutescens* (18) + *E. talpinus* (19); **Llg**, *L. lagurus*; **LSP**, *L. brandtii* (8) + *L. mandarinus* (9) + *L. gregalis* (10) + *L. raddei* (11); **MNS**, *M. miurus* (4) + *M. ochrogaster* (5) + *M. pennsylvanicus* (6) + *M. richardsoni* (7); **Ozb**, *Ondatra zibethicus*; **Psw**, *Prometheomys schaposchnikowi*; **OutG**, *N. mexicana* (34) + *C. brabensis* (35).

**Table 1.**
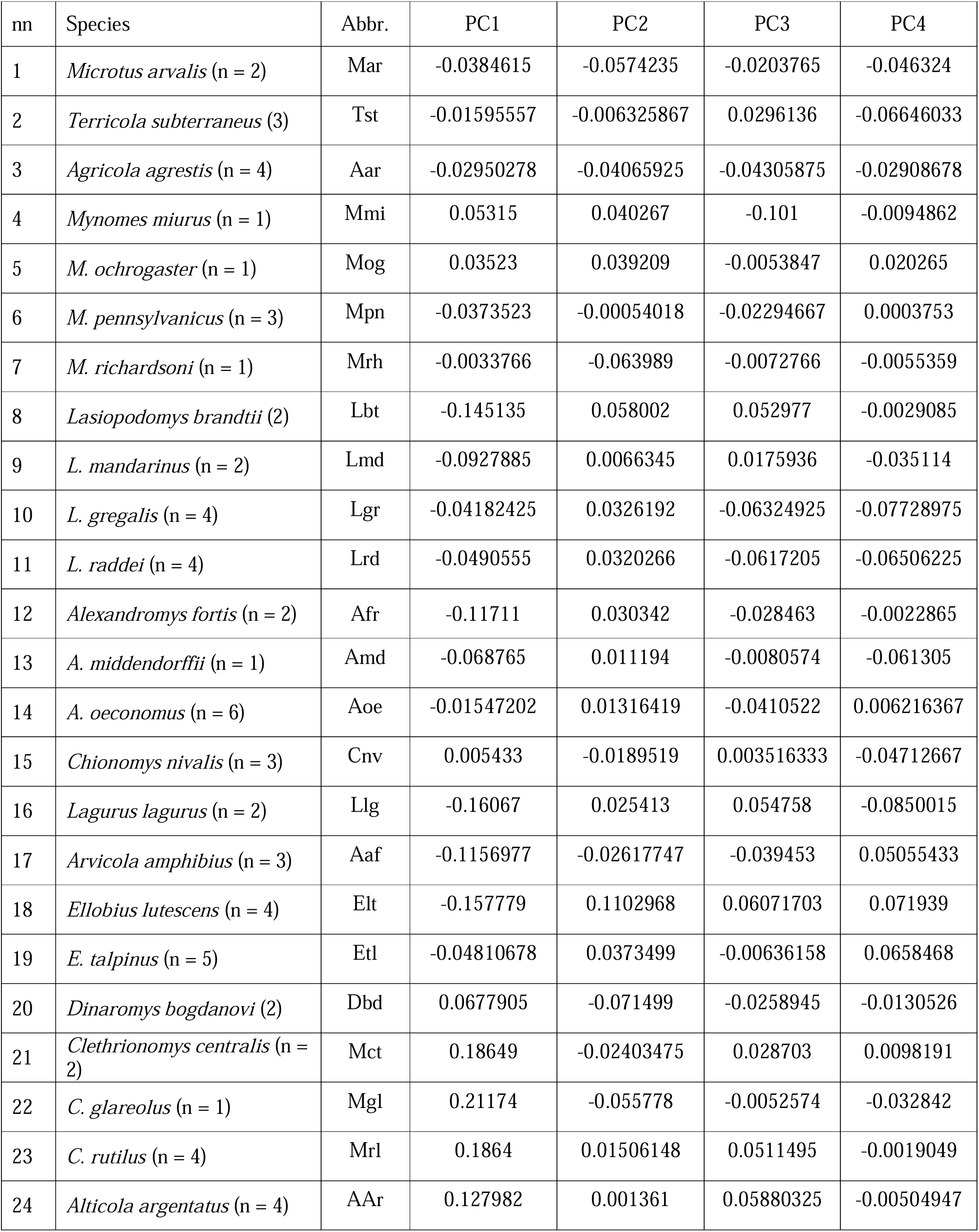
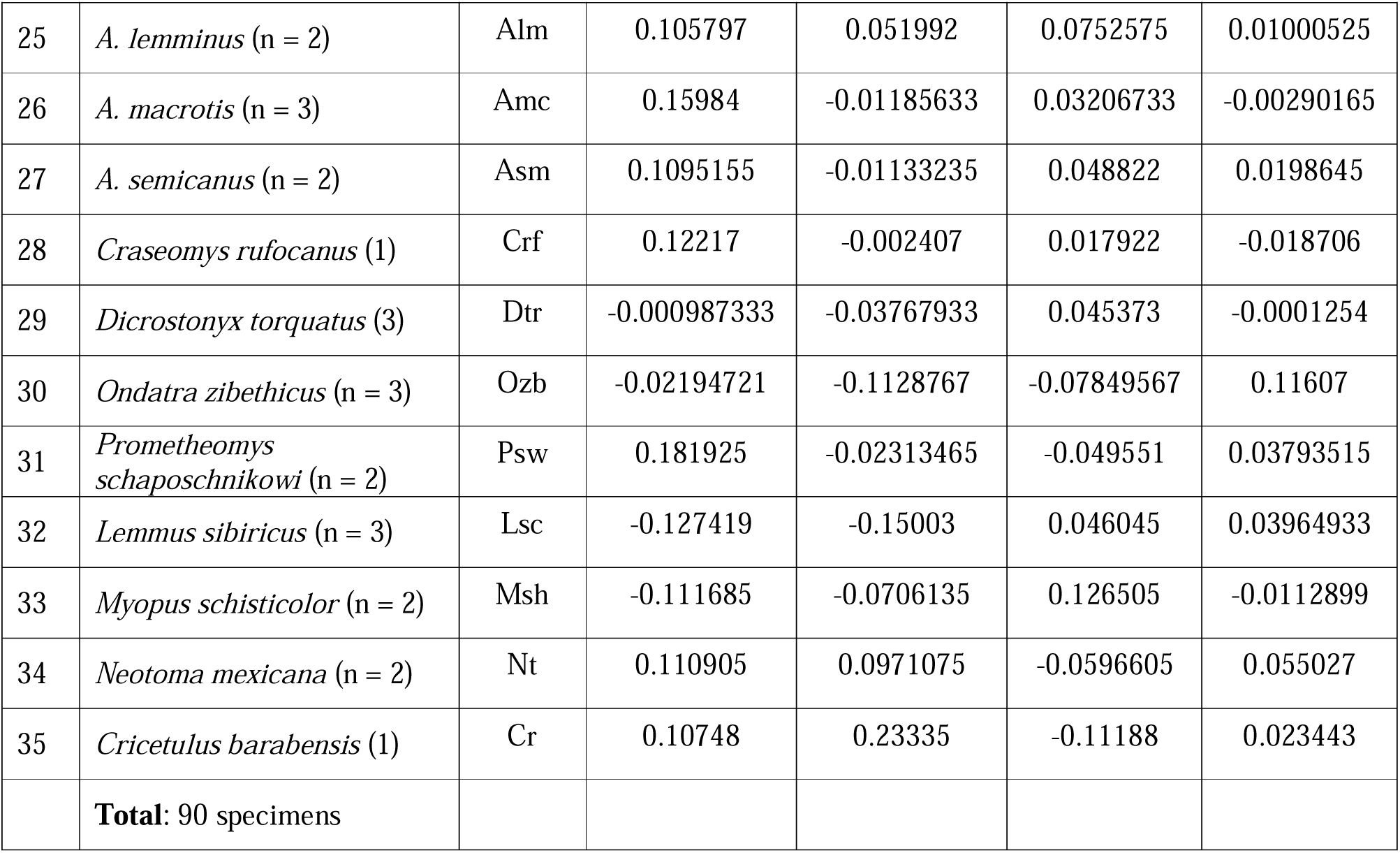
Mean values (centroids) of four principal components describing the shape variation of 35 species. The species order according to the mitogenomic tree species topology (Figure 1).

The next step was a description of the shape transformation along each axis in order to prepare the data for the RERconverge protocol (see the Introduction section). For this purpose, we defined six groups of paired shape descriptions, i.e., six “trait vectors.” The first trait vector (vI) describes the main shape variety along PC1. The vector is represented by a pair of reference shapes of *L. lagurus* (PC1 = −0.16) vs. *C. rutilus* (PC1 = 0.18). The second and third vectors describe the shape changes along the second principal component in relation to PC1’s variance: (vII) the pair of reference shapes of *L. sibirica* (the negative region of PC1; PC2 = −0.15) vs. *L. brandtii* (PC2 = 0.058); and (vIII) the pair of reference shapes of *D. bogdanovi* (the positive region of PC1; PC2 = −0.07) vs. *M. ochrogaster* (PC2 = 0.03) (Appendix 9A). The fourth vector (v-IV) describes the main shape disparity along PC3: the pair of reference shapes of *M. miurus* (PC3 = −0.10) vs. *M. schisticolor* (PC3 = 0.12). The fifth and sixth vectors describe shape alterations along the fourth principal component in relation to PC3’s variance: (v-V) the pair of reference shapes of *L. gregalis* (the negative region of PC3; PC4 = −0. 077) vs. *O. zibethicus* (PC4 = 0.11); (v-VI) the last pair of reference shapes of *L. lagurus* (the positive region of PC3; PC4 = −0.08) vs. *L. sibiricus* (PC4 = 0.03) (Appendix 9B).

We combined six pairs of shape comparisons and two additional species, *A. oeconomus* and *M. richardsoni*, into a “reduced subset” (see the Methods section) and repeated the PCA using the expanded landmark set of 121 Lms (Figure 2F). The analysis was performed on 12 species (*L. lagurus* and *L. sibiricus* participated in two pairs). The correctness of the results was checked by paired tests performed on two blocks of Procrustes distance values (Table 2). The tests in Table 2 support the applicability of our approach to visualization via PCA with the reduced subset.

**Table 2.** The Procrustes distance values (PrD, distance from the centroid position) of 12 reference species used for visualisation. The ’main set’ column contains the PrD calculated in the main PCA; the ’subset’ column contains the PrD obtained in the PCA calculated for visualisation (reduced subset). Statistics are shown at the bottom of the table. Tests are calculated using PAST.

| nn | Species | PrD: Main set (71 lms) | PrD: Subset (121 lms) |
| --- | --- | --- | --- |
| 1 | *M. richardsoni* | 0.10605 | 0.07628 |
| 2 | *A. oeconomus* | 0.10597 | 0.08069 |
| 3 | *L. gregalis* | 0.14406 | 0.12242 |
| 4 | *M. ochrogaster* | 0.12187 | 0.12342 |
| 5 | *L. brandtii* | 0.17369 | 0.13874 |
| 6 | *O. zibethicus* | 0.18782 | 0.14244 |
| 7 | *D. bogdanovi* | 0.13392 | 0.1436 |
| 8 | *M. miurus* | 0.14335 | 0.14854 |
| 9 | *L. sibiricus* | 0.17763 | 0.17016 |
| 10 | *L. lagurus* | 0.20028 | 0.17065 |
| 11 | *C. rutilus* | 0.19620 | 0.18896 |
| 12 | *M. schisticolor* | 0.24271 | 0.21652 |
|  |  | n = 12 | n = 12 |
|  | Mean: | 0.16113 | 0.14354 |
|  | Variance: | 0.0017 | 0.0016 |
| t-test = 1.0407, p-value = 0.30931 (9999 permut. runs: p = 0.3082)<br>F-test = 1.0724, p-value = 0.909 (corrected p = 0.907)<br>Mann-Whitney rank test: z = 0.8371, p-value = 0.4025 (corrected p = 0.409) |  |  |  |

The first vector describes four interpretable shape transformations (Figure 8A). Each transformation defines general disparities between two groups in terms of the first PCA (Figure 6): the first group consisted of the main arvicoline genera and the second group was composed of the “ancestral state” along PC1. The first group is represented by *L. lagurus*, and the second one by *C. rutilus*. Therefore, the “progressive state” of the PSP complex, which is derived from evolutionary features of the first group of arvicolines, is as follows (Figure 8A: light blue pattern):

**Figure 8.**
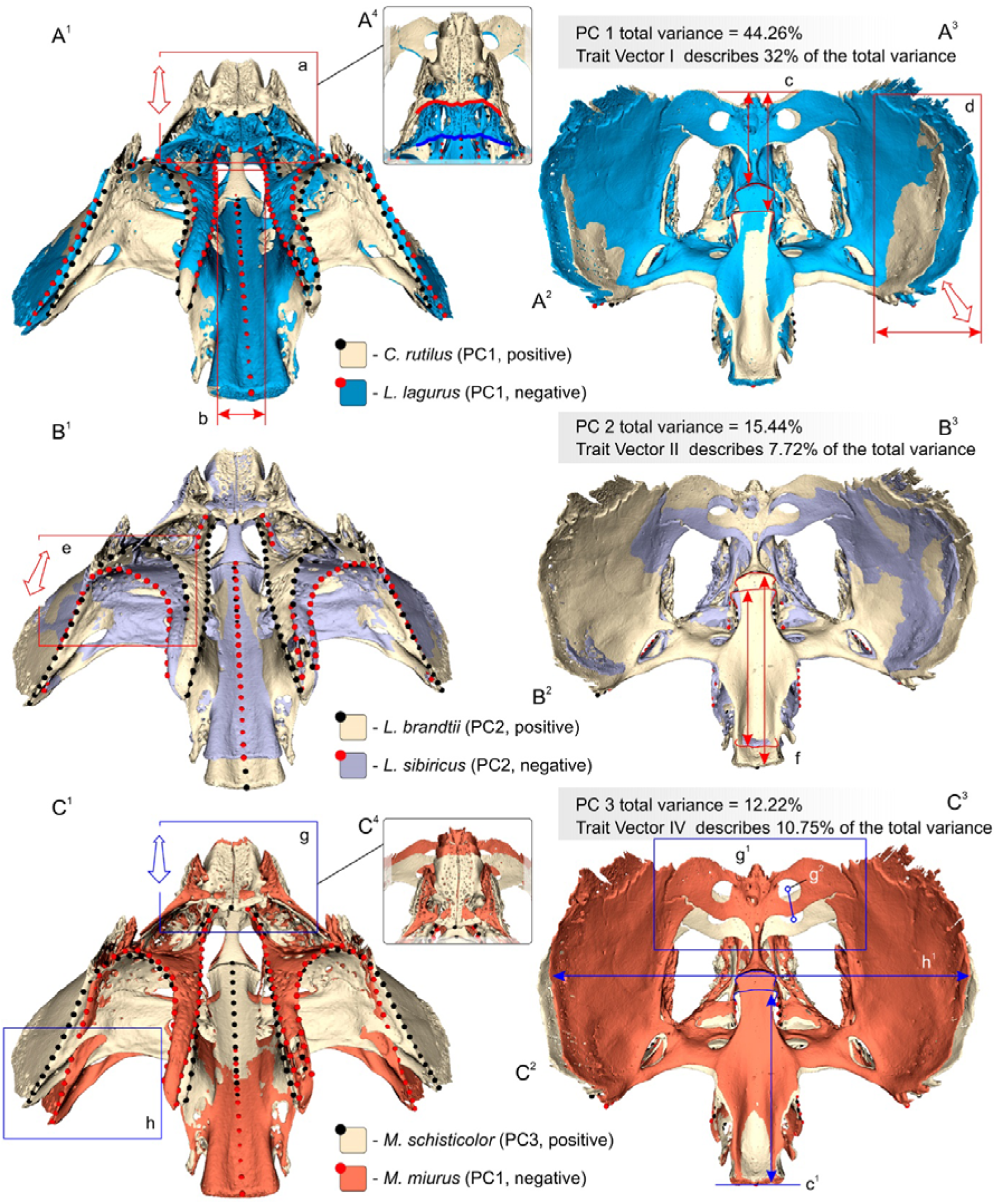
Results of the pairwise comparison of 3D models of the PSP complex according to the description of the shape transformations along the I, II and IV trait vectors (see Appendix 9). The comparisons were carried out using SlicerMorph (extension of 3D Slicer): each ’PCA Warped Model’ was saved separately; two such models were combined to obtain a final image using SlicerMorph again. (A^1^) Pairwise comparison of *C. rutilus* and *L. lagurus* (trait vector I); the PSP complex shown in ventral posterior aspect; (A^2^) same comparison, the complex shown in dorsal aspect (from the surface of the cranial cavity); (A^3^) plate with statistics: total variance of PC1 and variance absorbed by the first vector (see Appendix 11); (A^4^) enlarged palatine in ventral view showing differences between the shapes; (B^1^) Pairwise comparison of *L. brandtii* and *L. sibiricus* (trait vector II); the PSP complex shown in ventral posterior aspect; (B^2^) the complex shown in dorsal aspect; (B^3^) plate with statistics: total variance of PC2 and variance absorbed by the second vector; (C^1^) Pairwise comparison of *M. schisticolor* and *M. miurus* (trait vector IV); the PSP complex shown in ventral posterior aspect; (C^2^) the complex shown in dorsal aspect; (C^3^) plate with statistics: total variance of PC3 and variance absorbed by the fourth vector; (C^4^) enlarged palatine in ventral view showing differences between the shapes. Not to scale. See also appendix 10. *Key*: **a**, trait vector I [vI] — disparity in the horizontal lamina relative position; **b**, vI — disparity in the mesopterygoid fossa width; **c**, vI — disparity in the presphenoid/basisphenoid contact position (c^1^ similar transformation in the v-IV); **d**, vI — disparity in the sphenoid alae size; **e**, vII — disparity in the position of the anterior margin of the parapterygoid fossa; **f**, vII — disparity in the body length of the basisphenoid; **g**, v-IV — disparity in the position of the rostral parts of the palatine and presphenoid; **g^1^**, the transformation in the relative position of the orbital wing of the presphenoid; **g^2^**, the trajectory of the shift of the optic foramen (cf. the foramina in A^2^); **h**, v-IV — disparity in the width of the cranial cavity (shown for the ectopterygoid crest; h1, shown for the cranial cavity in the dorsal view).

(a) The horizontal lamina of the palatine as a whole has moved posteriorly (caudally) and superiorly (dorsally), absorbing anterior parts of the longitudinal laminae; at the same time, (b) the longitudinal laminae have moved closer together, resulting in a slight reduction in the width of the choanae. (c) The presphenoid/basisphenoid junction has moved anteriorly in contrast to the posterior movement of the horizontal lamina; i.e., the presphenoid has become short relative to the stable basisphenoid/basioccipital junction. (d) The overall size of the alisphenoid/orbitosphenoid (= the sphenoid ala) increased, and the presphenoid/sphenoid alae together have risen upward relative to the basisphenoid body. Besides, we can see the anterior displacement of the lateral processes of the basisphenoid. These four characteristics define the main differences between typical arvicolines and the Clethrionomyini/Prometheomyini group (and hamsters in a more global view): the “progressive” morphotype B2 (Figure 4) of the posterior palatal margin is formed by the significant posterodorsal shift of the horizontal lamina; therefore, clethrionomyins’ morphotype “A” (Figure 3) should be defined as “primitive” in the strict arvicoline trait space. The discrepancy between the “ancestral” position of the hamsters and their “quasi-progressive” morphotype “B1” (Appendix 3) can be reconciled as two evolutionary trajectories: “morphotype B1 → morphotype A” and alternatively “morphotype B1 → morphotype B2 [morphotype B3].”

Another feature distinguishing the two groups is the “visibility” of the presphenoid/basisphenoid junction above the posterior palatal margin: clethrionomyins, *Prometheomys*, and hamsters have a well-visible junction due to the long presphenoid body, whereas in typical arvicolines, the junction is hidden by the posterior margin of the palate. Nevertheless, this characteristic shows a large variation along PC1.

The upward rise of the presphenoidal orbital wings, together with the sphenoidal alae of typical arvicolines, is a consequence of the taxonomic group’s attempt to “move the brain away from high-crowned teeth.” In this context, it is quite obvious that the hypsodonty of Clethrionomyini has evolved in a different way as compared to typical arvicolines. and this hypothesis requires special investigation.

The second trait vector reflects shape changes within the first group of arvicolines (Figure 6) in a pairwise comparison of *L. brandtii* and *L. sibiricus* (Figure 8B). The vector describes two additional interpretable shape transformations that here explained the more specialized species *L. sibiricus* (pale violet pattern):

(e) The anterior edge of the parapterygoid fossa has shifted posteriorly and slightly superiorly with increasing pterygoid thickness. At the same time, the anterior end of the longitudinal lamina has shifted slightly superiorly, which, together with the fixed position of the horizontal lamina, represents the most progressive morphotype “B3” of the posterior palatal margin. The shift of the anterior margin of the parapterygoid fossa and the fixed position of the lateral process of the basisphenoid allow this transformation to be interpreted in the sense of Kesner’s statement about an increase in the anterior and medial vector components of the internal pterygoid muscle during graminivory development (Kesner 1980). Accordingly, the second vector revealed differences in the muscle vectors between *L. brandtii* and *L. sibiricus* and probably differences in graminivory specialization or in the structure of other components of a masticatory system. (f) There is a reduction in the length of the basisphenoid, topologically implying a shift of presphenoid/basisphenoid and basisphenoid/basioccipital joints. Next, we describe the fourth vector (see the next section for details).

The fourth trait vector reflects shape alterations along the third axis in the pairwise comparison of *M. schisticolor* and *M. miurus* (Figure 8C). The vector describes two further shape transformations, which are explained by *M. miurus* (orange pattern):

(g) The overall elongation of the perpendicular lamina of the palatine shifts the horizontal lamina of the palate and the anterior part of the presphenoid anteriorly together with the orbital wing. The significant shift of the orbital wing is marked by the shift of the optic foramen, not similarly to the presphenoid changes along PC1 (Figure 8: A(c)). Nonetheless, the main shape alterations along the fourth vector are associated with (h) the general narrowing of the cranial cavity in the PSP complex area, as evidenced by a decrease in its width between the sphenoid alae and between the ectopterygoid crests. Among arvicolines, there are two genera with well-pronounced morphological trajectories toward the “narrow-headed” morphotype, namely *Lasiopodomys* and *Mynomes*. Appendix 9 shows the differences in the direction between the trajectories; these dissimilarities are probably explained by a difference in the genotypic background between the taxonomic groups.

The third trait vector reflects shape changes within the “intermediate group” represented by *M. ochrogaster* and *D. bogdanovi*. The vector explains a low percentage variance (3.7%; see Appendix 10 and the Discussion section). Both species are quite similar, and the single characteristic reveals elongation of the posterior part of the palatal horizontal lamina in *M. ochrogaster*. This characteristic differs from trait “a” of the first trait vector (Figure 8A) by the fixed position of the anterior part of the palate and by the absence of a superior movement.

The fifth trait vector denotes shape changes in relatively specialized species *O. zibethicus* and *L. gregalis* (Appendix 10B) along PC4, which are in the “narrow-headed” range of PC3 (see description [g] above). The alterations explain the overall elongation of the PSP complex of *Ondatra* vis-à-vis the complex’s overall width that this similar between the two species. Here we mean “elongation” and “similar width” in relative terms because the size differences between these species are large. Nevertheless, *Ondatra* is classified here as a relatively narrow-headed species.

The last, sixth, trait vector reflects shape transformations along PC4 in the “broad-headed” region of PC3 between *L. sibiricus* and *L. lagurus* (Appendix 10C). *Lagurus* has a wider PSP complex and a more massive perpendicular lamina of the palate than does *Lemmus*.

## Discussion

### General remarks

The comparative morphological analysis based on the micro-CT data and specialized software applications (e.g., Avizo) enabled us to revise traits of the PSP complex, including key features of the palatine highly important for the definition of Arvicolinae taxonomic groups (Zazhigin 1980; Pozdnyakov 2008; Robovský et al. 2008; Abramson et al. 2020).

The exhaustive description of the palatine helped us to homologize the main morphological characteristics of Arvicolinae and to define some new ones. We defined the “pterygoidal process” of the palatal perpendicular lamina (Ch1), the “basisphenoidal process” of the perpendicular lamina (Ch2), and the “longitudinal lamina” of the palate with its “lamellar ridge” (Ch3, 4). These characteristics are described and illustrated for the first time for arvicolines and outgroup members: *Cricetulus* and *Neotoma* (Figures 3 and 4; Appendices 3, 4, and 6).

A special purpose of the current study was a revision of the characteristics of the palatal horizontal lamina owing to a specific morphotype of the palatine within the tribe Clethrionomyini. We were able to homologize the “major palatine foramen” (Ch5) and the “major palatine groove” (Ch6), which are related to the passage of the major palatine artery from the pterygopalatine fossa to the hard palate surface and next into the incisive foramen (Appendices 6 and 8). Homologies were established via original segmentation of our micro-CT data and with the help of detailed studies by Quay (1954), Evans (1993), and Wible (2008).

The next group of palatal features is related to the passage of the minor palatine artery. Here we were able to homologize the “palatine sulcus” (Ch7), which gives the artery access to the palatal space, and a “minor palatine groove” (Ch8). Among the examined species, the former characteristic was found only in cricetines (Appendices 4 and 8). A cricetine-like state of the minor palatine groove was detected in cricetines and *Prometheomys*. Homologies were established by means of the micro-CT data and detailed research of Fabre et al. (2018) on rodents and studies by Evans (1993) and Wible (2008) as a guide.

The “posterolateral palatal pit” (Ch9) was the most difficult to describe; it proved to be undeveloped in the outgroup species and usually hypertrophied among arvicolines.

The function of the posterolateral pit is still unknown. According to dissection of fluid-preserved specimens of *L. brandtii* (unpublished data), we revealed a large dense gland-like structure that completely fills the pit. It is likely that this entity is related to the soft palate salivary glands described for Muridae (e.g., Monroy et al. 2013), but we did not find specific studies on cricetids.

Furthermore, functional significance of the posterolateral palatal pit in relation to hypsodonty stages and/or the propolinal movement of the mandible in the sense of Kesner (1980) should be proposed. In the absence of published data on arvicolines, we used an existing description of oryzomine rodents (Weksler 2006).

Weksler’s phylogenetic analysis based on morphology included some important characteristics here that mark the expression of the posterolateral pit [character 34: four states from “pits absent” (0) to “pits always present as perforations within deeply recessed fossa” (3); (Weksler 2006: 35)], the depth of the parapterygoid fossa [character 35: three states from “fossa at the level of the palate” (0) to “fossa deeply excavated, reaching the level of the mesopterygoid roof” (2); *ibid*: 36], and the height of the molar crown and the type of occlusal surface [character 54: two states from "molars bunodont and brachydont" (0) to "molars planar and hypsodont" (1); *ibid*: 45].

Weksler’s analysis uncovered no relation between hypsodonty and the expression of the posterolateral pit within oryzomyines. That author’s data matrix (Weksler 2006: 20) contains only two hypsodont species: *Holochilus brasiliensis* and *H. chacarius*. As a consequence, *H. brasiliensis* was found to have no posterolateral pit (state 0), and *H. chacarius* to have a weakly developed pit (state 2). On the other hand, both *Holochilus* species have a maximum depth of the parapterygoid fossa (state 2). Some bunodont species, e.g., *Oryzomys polius*, have a deep arvicoline-like posterolateral pit (state 3) together with the maximally deep parapterygoid fossa (state 2). Because bunodonty restricts the propolinal movement of the mandible, the deep pit in *Oryzomys* does not seem to be related to the expression of horizontal vectors of the masticatory cycle.

If our initial hypothesis about the function of the posterolateral palatal pit means strengthened palatal structure, then Weksler’s analysis raises new questions that require research in both taxonomic groups.

Another result of our morphological description was the homology of components of the alisphenoid canal. Arvicolines differ from cricetines by simpler composition of the alisphenoid canal, where the trigeminal nerve and a few vascular branches emerge from the canal lumen without additional foramina (Appendix 6). On the contrary, in hamsters, the exit of the trigeminal nerve and of some branches of the IMA are separated: the lumen of the alisphenoid canal gives rise to several branches of the IMA (including the external ethmoidal, the external ophthalmic, and probably the minor palatine artery); the caudal "foramen ovale accessorius" (Ch10) in the sense of Wahlert (1974) gives rise to the trigeminal nerve (Appendices 7C and 8). Thus, voles and lemmings have a single lumen of the alisphenoid canal, whereas hamsters have two.

### The posterior palatal margin

Examination of palatal morphology revealed two main morphotypes in the combination “the rostral end of the lamellar ridge and the posterior palatal margin.” Morphotype “A” is unique to Clethrionomyini (Figure 3). Morphotype “B” is subdivided into “quasi-progressive” morphotype “B1” of cricetines (Appendix 3) and two successive states of “progressive” morphotypes “B2” in voles and “B3” in the lemmings. *Ellobius* specimens showed a mode of transformation from B2 to B3 in terms of the combination of the lamellar ridge and the posterior margin, depending on high inflation of the horizontal lamina of the palate.

Our findings lead us to the conclusion that morphotype B1 characterizes only brachyodont and mesodont cricetines and, despite the apparent similarity between B1 and B2 (Figure 4), has no direct relation with the evolution of the palatal margin in arvicolines (or we cannot see it yet). How the advancement of hypsodonty formed *Clethrionomys*-like morphotype “A” and *Microtus*-like morphotype “B2” is still unknown, but Repenning (1968) and Kesner (1980) are fairly confident in stating a direct relation between hypsodonty, bone transformations, and the reduction of the vertical vector component in pterygoid muscles of arvicolines.

On the other hand, our shape analysis detected a well-interpretable morphological trajectory of transformation of morphotype “A” to morphotype “B2” through a shift of the horizontal lamina posteriorly and superiorly, with absorption of the anterior part of the longitudinal lamina (Figure 8A). Thus, we know the mode of morphotype transformation, but we cannot say how many times this has occurred during evolution. Another question is why clethrionomyins are the only extant taxonomic group to have morphotype “A“ despite the differences in hypsodonty among its members: *Clethrionomys s. lato* vs. *Alticola s. lato*.

The answer may come from a combination of phenotypic and genotypic datasets and a search for adaptive signatures in genomic data (Christmas et al. 2023), using technical possibilities of reconstruction of ancestral phenotypic states by identification of "similar genetic changes that correspond to similar phenotypic changes may indicate a genotype-phenotype relationship" (Redlich et al. 2023).

### Trait vectors for use with R package RERconverge

The second part of the study—our morphometric analysis based on the 3D data—was primarily intended to prepare the morphological data for the evaluation of genotype–phenotype relationships in RERconverge (Kowalczyk et al. 2019; Redlich et al. 2023).

Results of the morphometric analysis allowed us to define six “trait vectors,“ each describing the main shape transformations along a given principal component. The first vector describes disparity along PC1 in a pairwise comparison of shape of *L. lagurus* vs. *C. rutilus*; the second and third vectors reflect disparity along PC2 in two types of shape comparison: *L. sibirica* vs. *L. brandtii* and *D. bogdanovi* vs. *M. ochrogaster*; the fourth vector describes the disparity along PC3 in the shape comparison of *M. miurus* vs. *M. schisticolor*; the fifth and sixth vectors reflect the disparity along PC4 in two types of shape comparison: *L. gregalis* vs. *O*. *zibethicus* and *L. lagurus* vs. *L. sibiricus* (Appendix 6).

The morphological data should be consistent and relatively easy to interpret. Nevertheless, Appendix 6 shows more than one trait vector used to describe the second and fourth principal components. Given that the choice of the number of vectors is subjective, six is not our final preferred number. We could choose more. In this case, one of important questions is the hierarchy of the vectors, i.e., which are the most important and which can be ignored.

To solve this problem, we propose to use a morphospace function for variance estimation [see “morphospace size estimation” in refs. (Voyta et al. 2021, 2023)]. This approach is illustrated in Appendix 11 and takes into account the ratio from the respective pairwise comparison of variance to the total variance of each axis (Appendix 2A). If we know coordinates of any pair of objects (Table 1), then we can calculate how much of the variance describes their relation in morphospace as a percentage. In this way, we evaluated the variance of each vector and ranked them in descending order of the percentage of variance explained.

It also follows that a given pair of comparisons describes only a part of the variance of each axis. Thus, the greater the number of comparison pairs, the more detailed is the shape description. In our opinion, however, a substantial increase in the number of vectors is not justified for the purpose of assessing phenotype–genotype relationships. In this sense, a subsequent increase in the detail of the description (its complexity) can be achieved iteratively.

The first, second, and fourth vectors explain 32.0, 7.72, and 10.75 percent of the variance along PC1, PC2, and PC3, respectively (Figure 8). We consider the variance along these vectors to be more meaningful than that of the remaining vectors (Appendix 10). Thus, the first three principal components (Table 1) can be utilized to identify genes and enriched pathways significantly associated with transformation of the PSP complex shape. According to the tested protocols, RERconverge allows to examine each component separately as a set of continuous variables (Kowalczyk et al. 2019; Christmas et al. 2023). The interpretable biological meaning of each component is described by the first, second, and fourth trait vectors (Figure 8). The remaining trait vectors, which explain a smaller percentage of variance, can be applied by separating values of a continuous variable of each component into a subset suitable for a local part of the morphospace (e.g., a smaller number of species).

In the case of more than one possible trait vector per principal component, we propose applying cluster analysis to correctly transform the set of continuous variables into a multi-categorical trait. This type of data can be investigated with the help of a special extension of the RERconverge protocol (Redlich et al. 2023).

Figure 9 outlines two possible strategies to divide clusters into two subsets: a binary characteristic (two main clusters) or multicategorical traits (three clusters). Table 3 consists of three types of data described by the first trait vector. This approach can be applied to PC2–PC4 (Table 1). The classic cluster analysis can be replaced by the K-means method.

**Figure 9.**
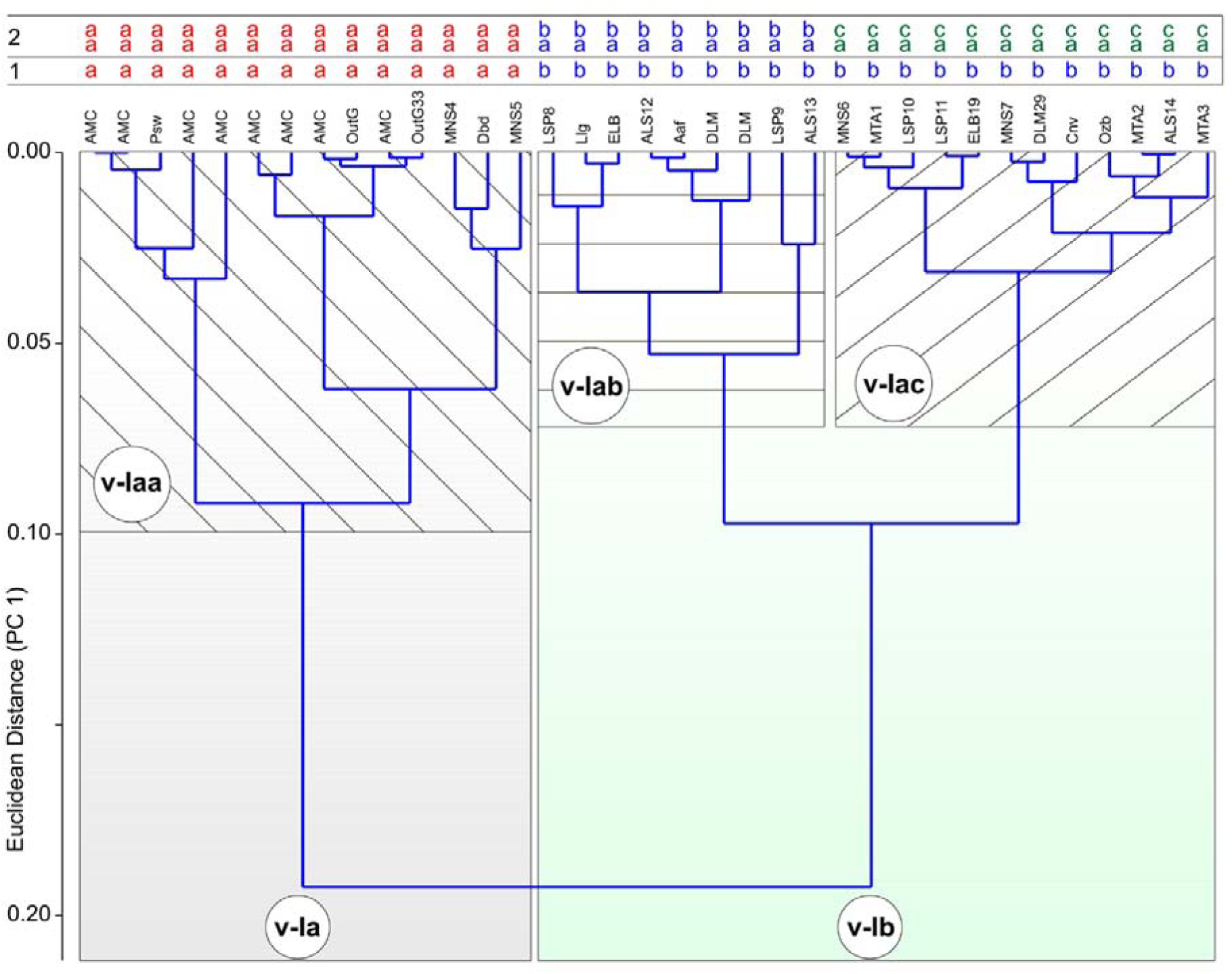
Results of cluster analysis (UPGMA; cophenetic correlation = 0.783) based on Euclidean distances of PC 1 values (Figure 6). *Key*: **1**, letter coding of the binary categorical trait (states ’a’, ’b’); **2**, letter coding of the multi-categorical trait (states ’aa’, ’ab’, ’ac’; used in the table 3). The first strategy of coding the categorical characters results in two clusters, which are used in the R package RERconverge as two states of binary character: **v-Ia**, cluster consists of outgroup + Clethrionomyini + *Prometheomys* + *Dinaromys* + *M. ochrogaster* + *M. miurus*; **v-Ib**, cluster consists of remaining groups of arvicoline. The second strategy of coding the categorical characters reveals three clusters, which are used in the R package RERconverge as three states of the multi-categorical character: **v-Iaa**, cluster is similar to the cluster v-Ia; **v-Iab**, cluster consists of *L. brandtii* + *L. lagurus* + *E. lutescens* + *A. fortis* + *A. amphibius* + *L. sibiricus* + *M. schisticolor* + *L. mandarinus* + *A. middendorffii*; **v-Iac**, cluster consists of the remaining groups of the arvicoline. PAST was used for cluster analysis.

**Table 3.** Three types of the codding of the first trait vector prepared for use in the R package RERconverge. Source of the trait vector values/states: ‘PC1 (vI)’ column contains continuous variables obtained in the PCA of the shape transformation (Figures 6, 7); ‘b-v-I’ column contains binary categorical states obtained in the cluster analysis on the base of the PC1 scores (Figure 9); ‘m-v-I’ column contains the multi-categorical states obtained in the same cluster analysis. *Key*: **Abbr**, acronyms of the species groups; **vI**, the first trait vector (Figure 8A).

| nn | Species | Abbr | PC1 (vI) | b-v-I | m-v-I |
| --- | --- | --- | --- | --- | --- |
| 1 | *Microtus arvalis* (n = 2) | MTA | -0.0384 | b | ac |
| 2 | *Terricola subterraneus* (3) | MTA | -0.0159 | b | ac |
| 3 | *Agricola agrestis* (n = 4) | MTA | -0.0295 | b | ac |
| 4 | *Mynomes miurus* (n = 1) | MNS | 0.0531 | a | aa |
| 5 | *M. ochrogaster* (n = 1) | MNS | 0.0352 | a | aa |
| 6 | *M. pennsylvanicus* (n = 3) | MNS | -0.0373 | b | ac |
| 7 | *M. richardsoni* (n = 1) | MNS | -0.0033 | b | ac |
| 8 | *Lasiopodomys brandtii* (2) | LSP | -0.1451 | b | ab |
| 9 | *L. mandarinus* (n = 2) | LSP | -0.0927 | b | ab |
| 10 | *L. gregalis* (n = 4) | LSP | -0.0418 | b | ac |
| 11 | *L. raddei* (n = 4) | LSP | -0.0490 | b | ac |
| 12 | *Alexandromys fortis* (n = 2) | ALS | -0.1171 | b | ab |
| 13 | *A. middendorffii* (n = 1) | ALS | -0.0687 | b | ab |
| 14 | *A. oeconomus* (n = 6) | ALS | -0.0154 | b | ac |
| 15 | *Chionomys nivalis* (n = 3) | Cnv | 0.0054 | b | ac |
| 16 | *Lagurus lagurus* (n = 2) | Llg | -0.1606 | b | ab |
| 17 | *Arvicola amphibius* (n = 3) | Aaf | -0.1156 | b | ab |
| 18 | *Ellobius lutescens* (n = 4) | ELB | -0.1577 | b | ab |
| 19 | *E. talpinus* (n = 5) | ELB | -0.0481 | b | ac |
| 20 | *Dinaromys bogdanovi* (2) | Dbd | 0.0677 | b | ac |
| 21 | *Clethrionomys centralis* (n = 2) | AMC | 0.1864 | b | ac |
| 22 | *C. glareolus* (n = 1) | AMC | 0.2117 | b | ac |
| 23 | *C. rutilus* (n = 4) | AMC | 0.1864 | b | ac |
| 24 | *Alticola argentatus* (n = 4) | AMC | 0.1279 | b | ac |
| 25 | *A. lemminus* (n = 2) | AMC | 0.1057 | b | ac |
| 26 | *A. macrotis* (n = 3) | AMC | 0.1598 | b | ac |
| 27 | *A. semicanus* (n = 2) | AMC | 0.1095 | b | ac |
| 28 | *Craseomys rufocanus* (1) | AMC | 0.1221 | b | ac |
| 29 | *Dicrostonyx torquatus* (3) | DLM | -0.0009 | b | ac |
| 30 | *Ondatra zibethicus* (n = 3) | Ozb | -0.02194 | b | ac |
| 31 | *Prometheomys schaposchnikowi* (n = 2) | Psw | 0.1819 | b | ac |
| 32 | *Lemmus sibiricus* (n = 3) | DLM | -0.1274 | b | ab |
| 33 | *Myopus schisticolor* (n = 2) | DLM | -0.1116 | b | ab |
| 34 | *Neotoma mexicana* (n = 2) | OutG | 0.1109 | b | ac |
| 35 | *Cricetulus barabensis* (1) | OutG | 0.1074 | b | ac |

### Essential shape comments

In the final part of the discussion, we provide a purely biological interpretation of the analysis of PSP complex shape, using the morphospace between PC1 and PC3 and the time of divergence events from a time-calibrated phylogenetic tree of Abramson et al. (2021).

The two most highly loaded trait vectors, the first one (32%, see Figure 8A3) and the third one (10.75%), reflect three most significant shape modifications that strictly define the relative position of arvicoline taxonomic groups in the morphospace with respect to three successive waves of adaptive radiation (Abramson et al. 2021). Members of two separate clades (*Lemmus* and *Myopus*) and (*Prometheomys* (*Ondatra* (*Dicrostonyx*))) represent the first wave of radiation (R1) and are widely distributed in the morphospace. The convex hulls of lemmings and musk rats occupy the morphospace together with arvicolines of the third radiation wave, whereas only *Prometheomys* is found in the morphospace together with Clethrionomyini. The latter represents the second radiation wave (R2). Convex hulls of the second and third waves (R3) do not overlap (Figure 10A).

**Figure 10.**
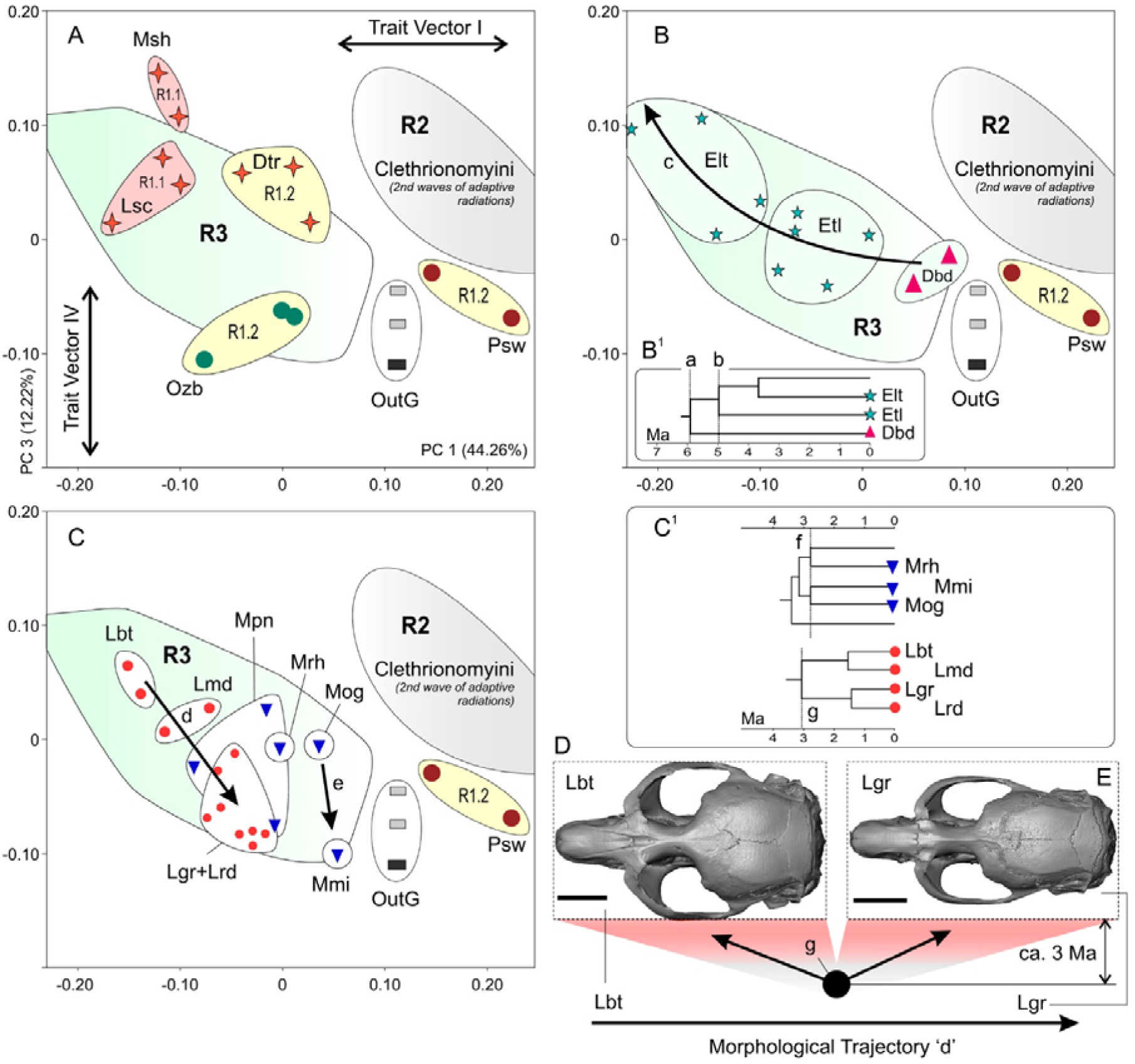
Interpretation of the Arvicolinae groups position within morphospace (PC1/PC3). (A) General scheme of morphospace relationships between three successive waves of the adaptive radiation of Arvicolinae as revealed by the mitochondrial genome (Figure 1); (B) relative position of members of the clade (*Dinaromys* (*Ellobius talpinus* (*E. lutescens* + *E. fuscocapillus*))); (B1) enlarged part of the time-calibrated phylogenetic tree (Abramson et al. 2021) with two evolutionary events marked as starting points of PSP complex shape divergence; (C) relative position of members of two clades: *Mynomes* spp. and *Lasiopodomys* spp. (both clades have extremely narrow-headed members); (C1) enlarged part of the time-calibrated phylogenetic tree with marking of two evolutionary events within the *Mynomes* and *Lasiopodomys* clades as starting points of the PSP complex shape divergence; (D) skull of an adult specimen of *L. brandtii* (ZIN 82114, ad2) in dorsal view; (E) skull of mature specimen of *L. gregalis* (ZIN 103094, ad2) in dorsal view. E and D are linked to form a pictogram showing the point of the evolutionary event occurred about 3 Ma ago at the divergence of the two species. Scale bars are 5 mm. *Key*: **a**, evolutionary event occurred about 6 Ma ago at the divergence of early representatives of the modern *Dinaromys* and *Ellobius*; **b**, evolutionary event occurred about 5 Ma ago at the divergence within the Ellobiusini; **c**, morphological trajectory of the shape transformations between evolutionary events ’a’, ’b’ and the recent state; **d**, morphological trajectory of the shape transformations between the states ’Lbt’ and ’Lgr’ during the period of about 3 Ma (event ’g’); **e**, morphological trajectory of the shape transformations between the states ’Mog’ and ’Mmi’ during the period of about 2.8 Ma (event ’f’); **f**, evolutionary event occurred about 2.8 Ma ago at the divergence within the *Mynomes*; **g**, evolutionary event occurred about 3 Ma ago at the divergence within the *Lasiopodomys*; **Dbd**, *D. bogdanovi*; **Dtr**, *D. torquatus*; **Elt**, *E. lutescens*; **Etl**, *E. talpinus*; **Lbt**, *L. brandtii*; **Lgr**, *L. gregalis*; **Lmd**, *L. mandarinus*; **Lrd**, *L. raddei*; **Lsc**, *L. sibiricus*; **Mmi**, *M. miurus*; **Mog**, *M. ochrogaster*; **Mpn**, *M. pennsylvanicus*; **Mrh**, *M. richardsoni*; **Msh**, *M. schisticolor*; **OutG**, outgroup; **Ozb**, *O. zibethicus*; **Psw**, *P. schaposchnikowi*; **R1.1**, first wave of the adaptive radiation, Lemmini clade; **R1.2**, first wave of the adaptive radiation, clade (*Prometheomys* (*Ondatra* (Dicrostonychini))); **R2**, second wave of the adaptive radiation, Clethrionomyini clade; **R3**, third wave of the adaptive radiation.

It is noteworthy that the shape of the PSP complex of *L. sibiricus* lies strictly within the convex hulls of R3. This position with respect to the first and third trait vectors can be interpreted as a similarity to R3 members in the topology of the palate (Figure 8: A(a)), in the length of the presphenoid (Figure 8: A(c)), and in the broad relative position of the sphenoid alae, denoting a “broad-headed” morphotype (Figure 8: C(h)). Other members of the R1 wave, with the exception of *Prometheomys*, are more or less divergent from the convex hulls of R3. *Prometheomys* is a more “narrow-headed” genus compared to R2 members. *Ondatra* tends to have “narrow-headed” morphology similar to that of R3 members.

The most unusual taxon of Arvicolinae, the Ellobiusini tribe, together with the “living fossil” vole *Dinaromys*, form a sister branch toward other R3 clades. We highlighted their positions in the morphospace and collated them with time points of their divergence (Figure 10: B, B1). Morphological trajectory “c” shows the supposed successive shape transformations between two evolutionary events and the most recent state. The first event (a) occurred ∼6 million years ago (Mya) during the divergence of early representatives of modern *Dinaromys* and *Ellobius*. The second event (b) took place ∼5 Mya during internal divergence of early representatives of the modern *E. talpinus*-and-*E. lutescens* clade. Thus, between 6 and 5 Mya, the PSP shape of the “*Dinaromys*-like” morphotype started to change to the “*Ellobius*-like” morphotype.

We should digress here to mention new information on the aforementioned relationship between *Dinaromys* and *Ellobius*, as revealed by an analysis of the nuclear genome recently obtained by Abramson’s team. The new phylogenetic information will be published soon (Abramson et al., *in press*) and involves a new topology of the phylogenetic tree, where *Dinaromys* is clustered with Lagurini and is located at a more basal position as compared to the Ellobiusini clade. On the one hand, these new relationships require a revision of morphological trajectory “c” because it must explain relationships *Dinaromys*/*Lagurus*. On the other hand, the new position of Ellobiusini does not invalidate trajectory “c” because ancestors of mole voles had a generalized arvicoline morphotype anyway, e.g., a *“Dinaromys*-like” morphotype. If we take into account the nuclear genomic data, then we should only add another trajectory *Dinaromys*/*Lagurus*, without discarding trajectory *Dinaromys*/*Ellobius*, because *Dinaromys* occupies the region of the “ancestral state” (Figure 6), and it is quite suitable for the role of the ancestral morphotype toward early representatives of Lagurini and Ellobiusini.

Another notable tendency in the arvicoline cranial shape is related to the existence of “narrow-headed” species. The best known species of this type is the narrow-headed vole *L. gregalis*. Another is the singing vole *M. miurus*. The former belongs to the *Lasiopodomys* clade. The latter species is affiliated with the *Mynomes* clade (Abramson et al. 2021). We also highlighted their positions in the morphospace and indicated the corresponding parts of the phylogenetic tree (Figure 10: C, C1). Morphological trajectories (d, e) of the two clades are significantly diverged along PC1, which in terms of trait vectors implies a greater similarity of the *Lasiopodomys* clade to R3 members, whereas the *Mynomes* clade tends toward R2 members. In this case, the vectors describe the topology of the palate (Figure 8: A(a)) and the length of the presphenoid (Figure 8: A(c)).

The presumed successive shape transformation along the morphological trajectories was determined for two evolutionary events. The first event (f) occurred ∼2.8 Mya during the divergence of early representatives of modern *M. miurus* and *M. ochrogaster*. The second event (g) took place approximately 3 Mya during the divergence of early representatives of modern *L. brandtii*/*L. mandarinus* and *L. gregalis*/*L. raddei*. Therefore, the “narrow-headed” morphotype of the Old World *Lasiopodomys* clade originated ∼3 Mya. The “narrow-headed” morphotype of the New World *Mynomes* clade arose independently at approximately the same time.

In conclusion, on the basis of the above findings, we are ready to formulate some questions about genetic datasets regarding the existence of technical possibilities for reconstruction of ancestral phenotypic states:

(i) Which genes are associated with the PSP complex shape convergence within the two main groups of arvicolines, R2 and R3?
(ii) Which genes are connected to the PSP complex shape convergence between *Prometheomys* and members of R2?
(iii) Are there differences in genes or their states between Ellobiusini and members of R3?
(iv) Which genes or their states determine morphological trajectory “c”?
(v) Are there similarities between the *Lasiopodomys* and *Mynomes* clades in the genes that determine the emergence of the “narrow-headed” morphotype?

## Conclusions

The current paper offers a protocol for the preparation of morphological data for the evaluation of genotype–phenotype relationships using R package RERconverge (Kowalczyk et al. 2019; Christmas et al. 2023; Redlich et al. 2023). Our morphospace analysis of the shape of the PSP complex in arvicolines revealed six trait vectors describing shape transformations along the first four principal components.

Our other protocol of morphometric data processing involves cluster analysis. This approach makes it possible to transform a continuous variable into a categorical one that remains suitable for the RERconverge calculations.

The obtained 3D dataset based on 71 landmarks can also be employed in another protocol, for the evaluation of phylogenetic signals in R package Geomorph (Adams 2014; Terray et al. 2022; Voet et al. 2022).

Biological significance of the detected morphological alterations was determined via interpretation of intergroup relationships within the morphospace along PC1 and PC3. The interpretations were based on two most highly loaded trait vectors—the first one (32%) and the third one (10.75%)—which together reflect three most significant shape modifications of the PSP complex that strictly determine relative positions of arvicoline groups in the morphospace with respect to three successive waves of adaptive radiation in the sense of Abramson et al. (2021).

One of important results is a revision of characteristics of the PSP complex, including key features of the palatine that are crucial for the definition of Arvicolinae taxa by means of the micro-CT data and specialized digital applications (e.g., Avizo).

This paper is the first contribution to the comprehensive analysis of the complicated evolution of cranial and mandibular parts connected by pterygoid muscles, as part of a more global investigation into adaptive evolution of Arvicolinae. A clue to this global puzzle lies in a search for closely correlated cranial and mandibular elements, so-called “modules” in the sense of Klingenberg (2009), which enable an analysis of a jaw fragment and assessment of changes in the skull. Repenning (1968) and Kesner (1980) wrote about the existence of such links. Research along this avenue will help to describe ancestral states of cranial elements (e.g., the PSP complex) based on fossil jaw fragments, which are much more often found in the fossil record then unique cranial fragments or whole skulls.

## Supplementary Materials

The following supporting information can be downloaded at: https://github.com/ZaTaxon/Evolution-of-the-skull-in-arvicoline-cricetids-Rodentia-.git.

Supplementary Material include **Table S1**, Information on analyzed samples (3D model dataset); **Table S2**, The list of 3D models used in the study and technical characteristics of micro-CT scanning (NeoScan N80 [FP]); **Figures S1–35**, Relative age groups of the analyzed specimens; **Figures S36–63**, Screenshots of the PSP complex with landmark positions in analyzed specimens. 3D model available via link: https://zin.ru/labs/evolgenome/morphology/index_Part1.html

## Acknowledgements

This study was funded by Project No. 19-74-20110 of the Russian Scientific Foundation. The study involved the collection materials at the Zoological Institute of the Russian Academy of Sciences (St. Petersburg; http://www.ckp-rf.ru/usu/73561/). The authors would like to thank their colleagues Vladimir S. Zazhigin, Maxim V. Sinitsa, Fedor N. Golenishchev, Natalia I. Abramson, Semyon Yu. Bodrov, Tatyana V. Petrova, Elizaveta Skalon, and Valentina A. Panitsina for constant discussions of various parts of the study and assistance with the conduct thereof. The authors are also grateful to Vladimir V. Platonov, Olga V. Makarova, and Eugenia R. Maksimova for technical help during the current project.

## Authors’ contributions

LV: conception, methodology, morphometry, statistical analysis, taxonomy and nomenclature, micro-CT data processing-segmentation, graphics, writing-original draft, and correspondence. DM: micro-CT and preliminary preparation of the image stacks of each analyzed specimen.

## Appendix 1

Screenshot of the PSP complex from 3D Slicer in ventral posterior view of specimen ZIN 102929 of *Craseomys rufocanus* with landmark positions (A^1^), and screenshots of the 3D cranial model of *C. rufocanus* (A^2-4^) and *Clethrionomys glareolus* (B^1–3^, ZIN 106573). (A^2^, B^1^) Skulls in lateral view; (A^3^, B^2^) skulls in dorsal view; (A^4^, B^3^) Lateral radiograph of molar roots; (A^5^) photo of the original label of *C. rufocanus*. Scale bar is 5 mm. Both specimens are shown in the supplementary material (Figures S11 and S28). The local age scale was filled by comparing age-related characters such as the lateral contour of the skull, temporal line expression and root development (for root-bearing species only). See remark below.

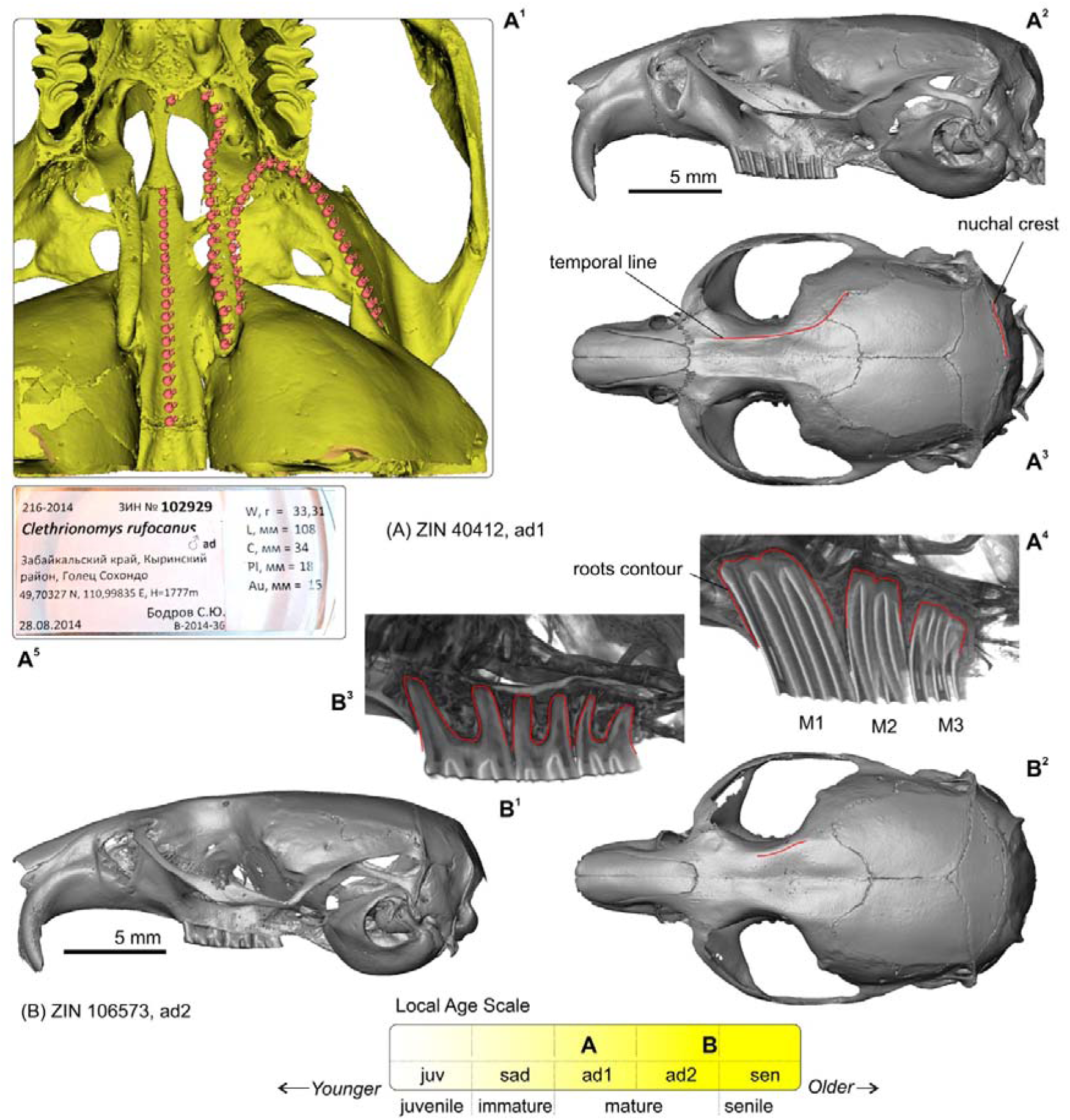

### Remark

The first specimen of *C. rufocanus* shows relatively strong development of the temporal and nuchal ridges in contrast to a weak root formation (the initial stage). On the other hand, the second specimen of *C. glareolus* has a very weakly developed braincase sculpture with strongly developed tooth roots. This feature incongruence is explained by interspecies differences in hypsodonty: *C. rufocanus* is more hypsodont species than *C. glareolus*. The correct age determinant should make use of several features, not only root development.

## Appendix 2

Result of two tests to determine the number of ’biologically loaded’ principal components. (A) Brocken-stick model plot calculated by default in PAST based on PCA analysis of PSP complex shape variation; (B) special use of Horn’s scree plot based on combining structured (PSP complex shape) and random data sets. The Principal Component Analysis statistics are shown in the two tables: the top table represents the structured data; the bottom table represents the random data (tables limited by the 15th axis).

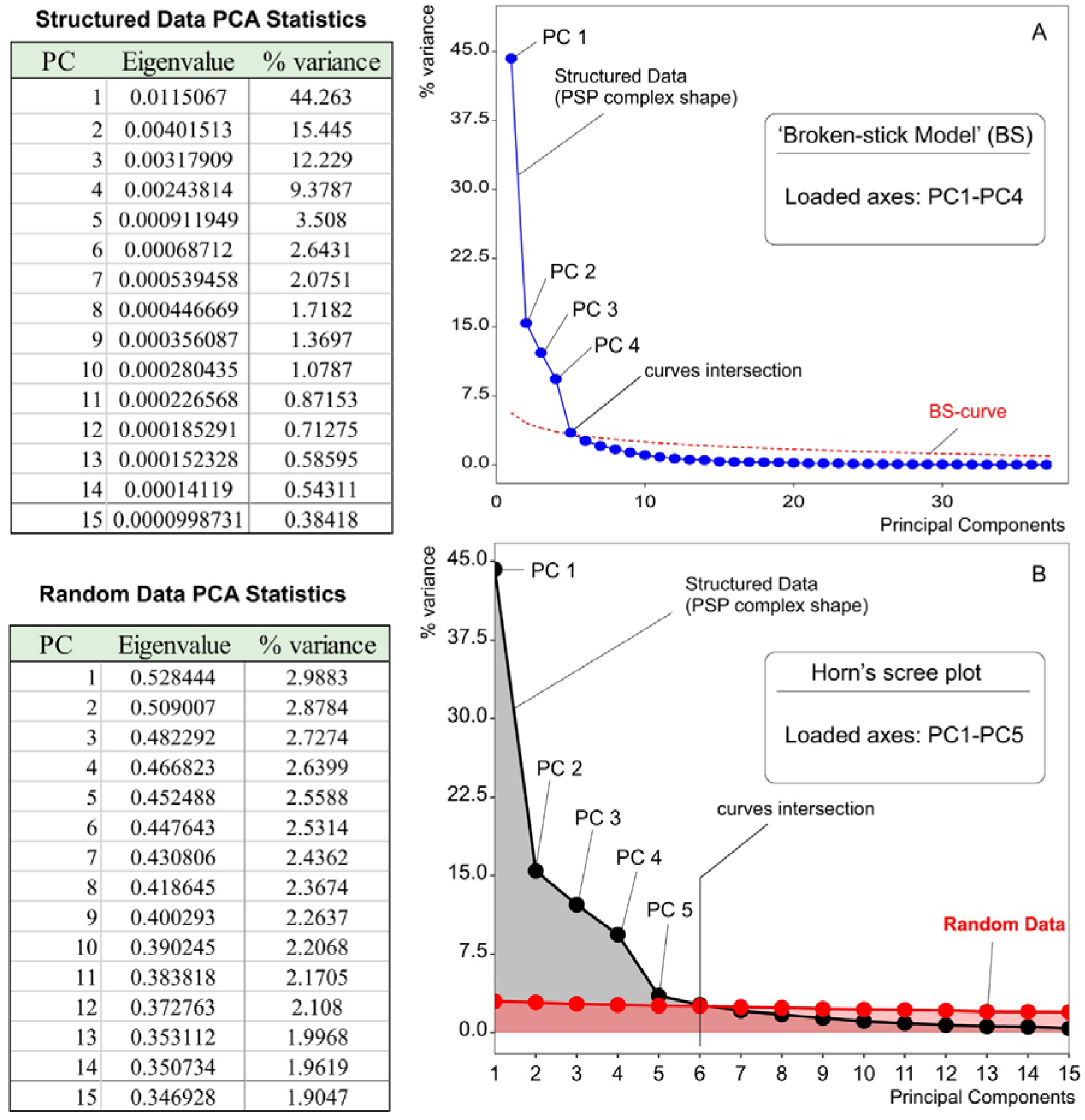

## Appendix 3

Isolated palatine of hamster, *Cricetulus barabensis* (ZIN 101697; outgroup). (A) Isolated palatine in ventral view; (B) isolated palatine in dorsal view; (C) right palatine bone with surrounding bones (grey) in medial aspect; (D) left palatine with surrounding bones (grey) in ventral lateral aspect (fossae of external and internal pterygoid muscles origin shown); (E) pictogram of morphotype B. *Key*: **a**, micro-CT artefacts; **bs**, basisphenoid; **epf**, external pterygoid fossa; **ham**, pterygoid hamulus; **lng**, longitudinal lamina (lamellar ridge); **pal-pm**, palatal posterior margin; **ps**, presphenoid (basi- and presphenoid contact marked by red line). Arrow marks the end of the lng at the contact with the pterygoid.

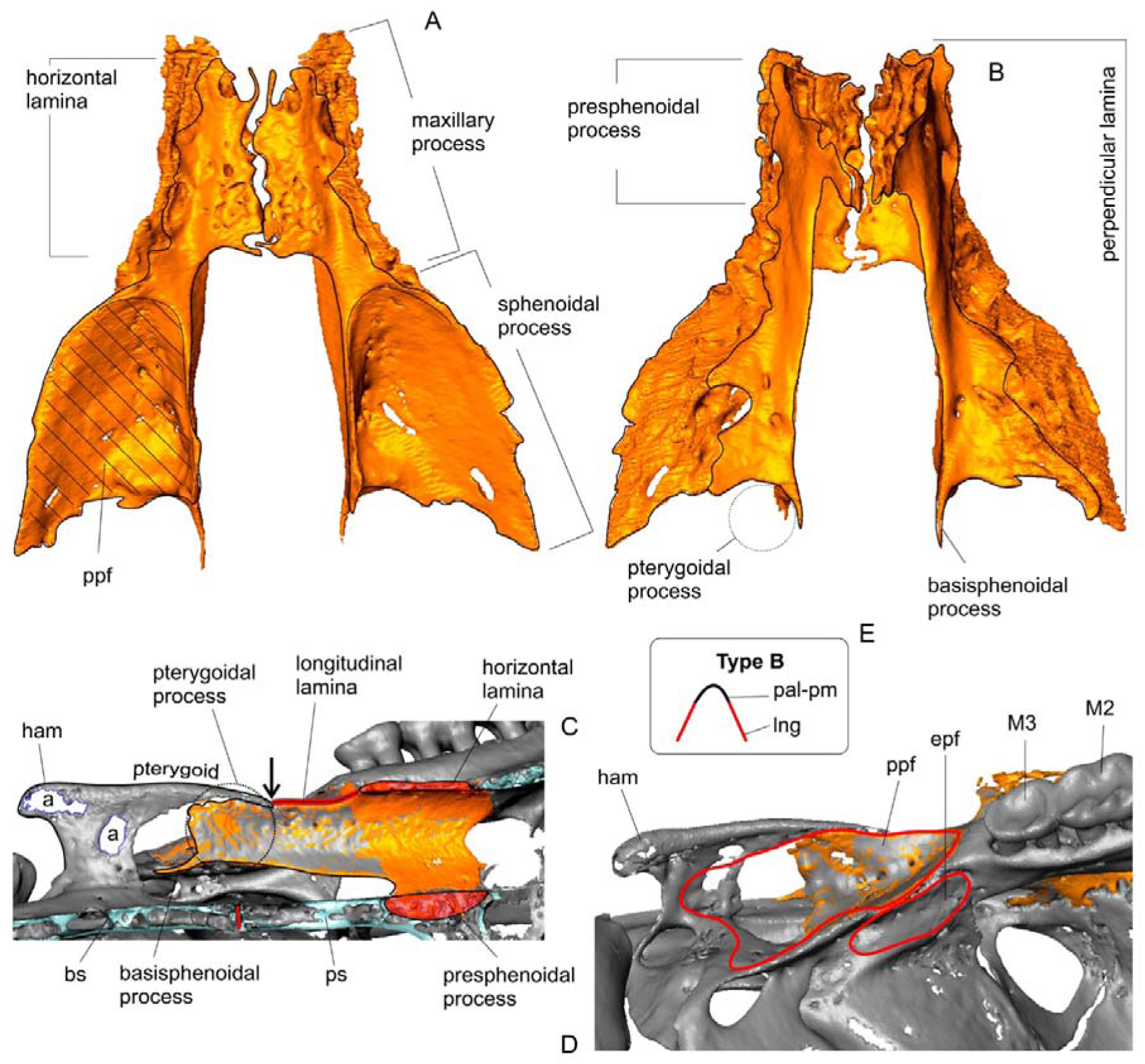

## Appendix 4

Position and proportion of the parapterygoid fossa in Cricetinae and Arvicolinae. (A) Cropped image of the skull of *Cricetulus barabensis* (ZIN 101697) in ventral view; (A1) explanatory drawing; (B) cropped image of the skull of *Alticola argentatus* (ZIN 77489) in ventral view; (B1) explanatory drawing; (C1) diagram showing the topological relationship between the ventrolateral ridge (vlr) and the ridge of the longitudinal lamina (lng) in *C. barabensis* (true for *Neotoma*); (C2) *ibid.*, *A. argentatus* (true for arvicolines); (D) pictograms of two morphotypes showing the relationship between the posterior palatal margin (pal-mg) and lng. Not scaled.

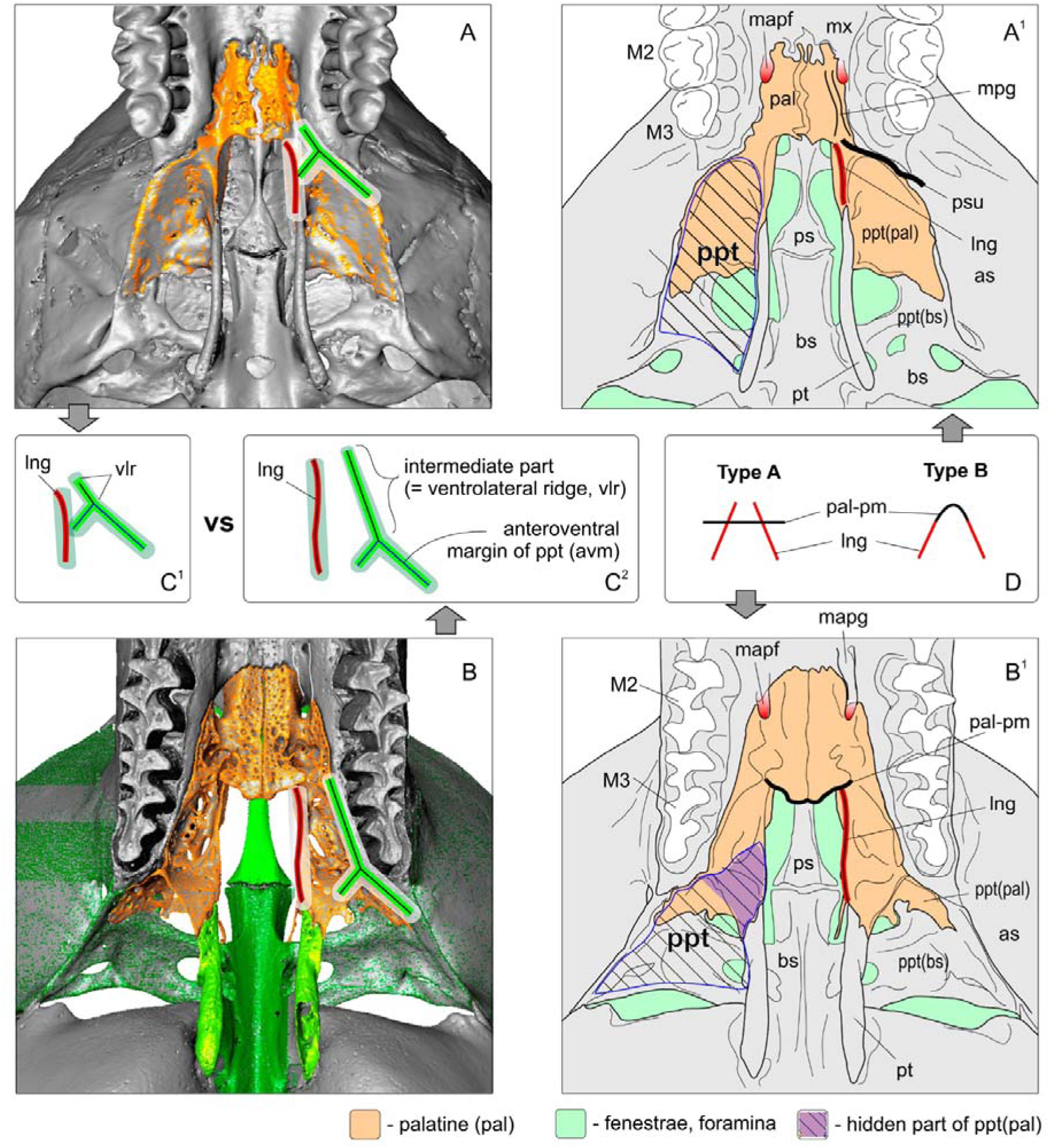

*Key*: **as**, alisphenoid; **avm**, anteroventral margin of ppt; **lng**, longitudinal lamina (lamellar ridge); **mapf**, major palatine foramen; **mapg**, major palatine groove; **mpg**, minor palatine groove; **mx**, maxilla; **pal**, palatine; **pal-pm**, posterior palatal margin; **ppt**, parapterygoid fossa; **ppt(bs)**, basisphenoid portion of ppt; **ppt(pal)**, palatal portion of ppt; **ps**, presphenoid; **psu**, groove/canal of the minor palatine artery; **pt**, pterygoid; **vlr**, ventrolateral ridge of the perpendicular lamina.

## Appendix 5

Position and development of the posterior nasal spine (pns) of the palatine of *Ellobius talpinus* (ZIN 82955). (A) Skull in dorsal view, the clipping plane is shown; (B) skull in lateral view; (C) enlarged part of the skull, in ventral lateral aspect; (D) enlarged part of the skull, in ventral posterior aspect; (E) enlarged part of the skull, in ventral aspect. No scaled.

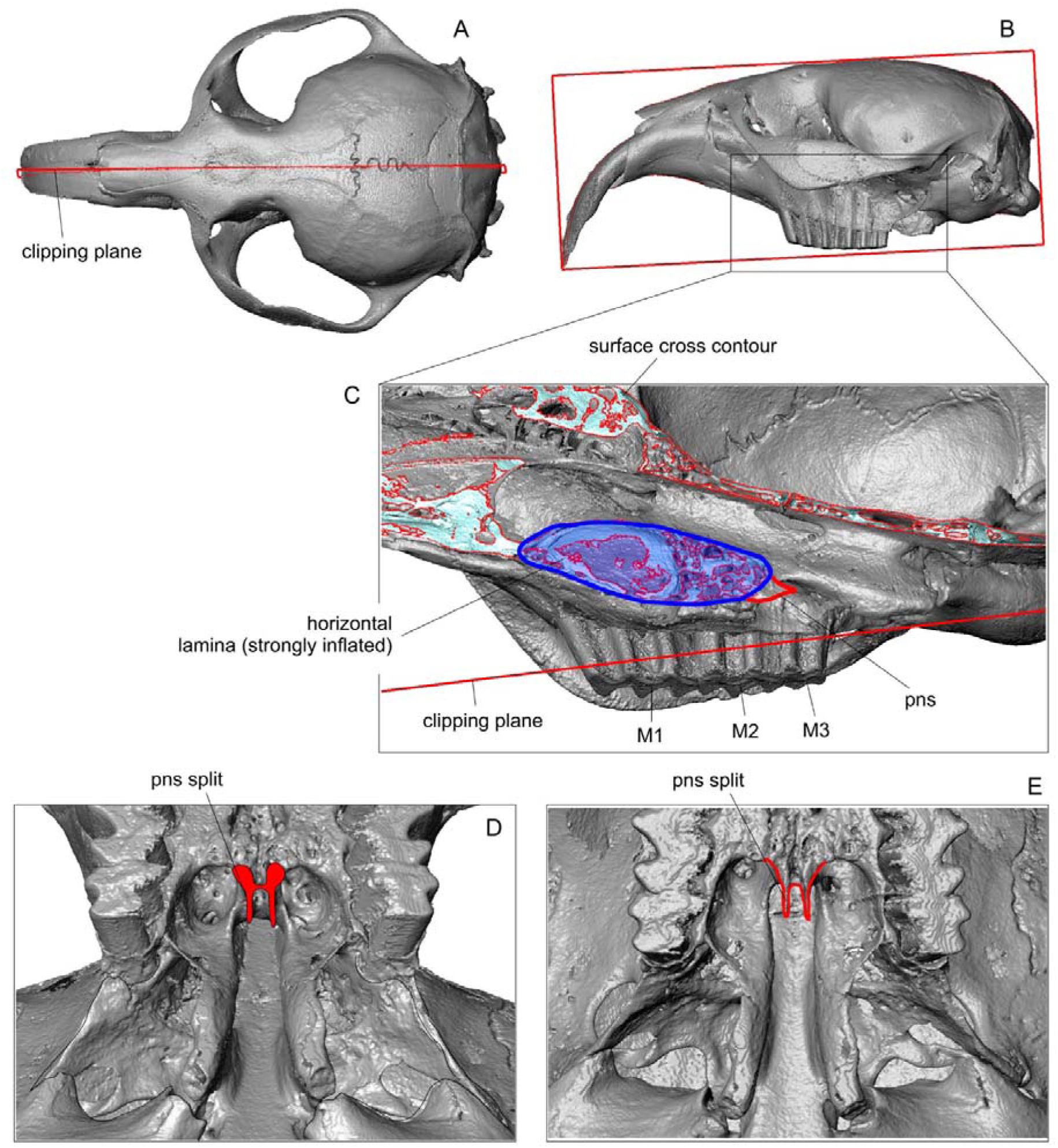

## Appendix 6

Reconstruction of the major palatine artery passage (detailed Avizo segmentation) of *Alticola argentatus* (ZIN 77489) and presumed branching of the internal maxillary artery, obtained from the groove casts of the internal surface of the braincase. Topology of the vessels according to Brylski (1991) and Evans (1993).

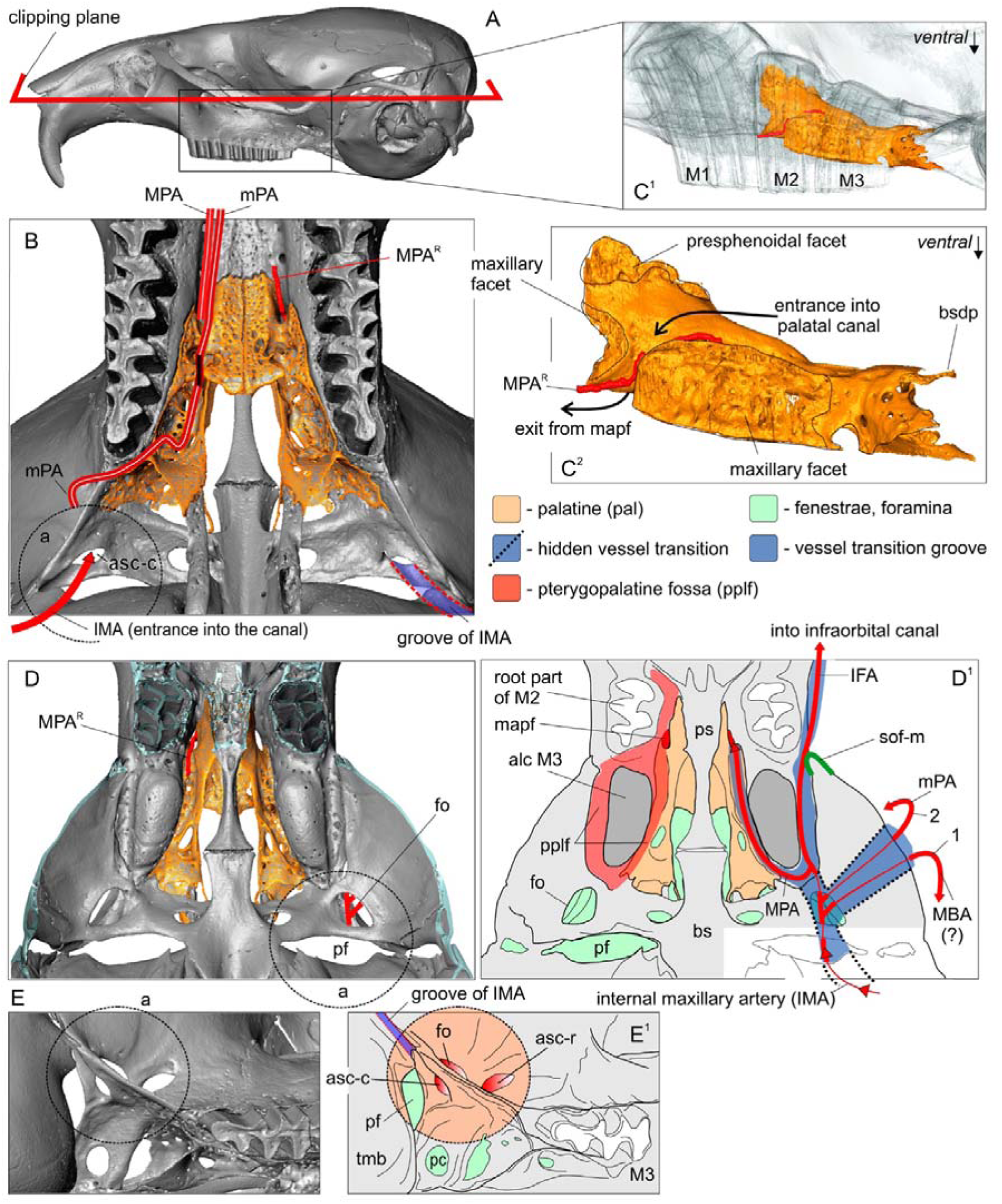

(A) lateral view of skull; (B) enlarged part of the skull in ventral view; (C^1^) transparent lateral view of skull with coloured palatine; (C^2^) isolated palatine in ventral view with reconstructed position of the major palatine artery; (D) enlarged part of the skull, view from the internal surface of the braincase; (D^1^) explanatory drawing; (E) enlarged the right alisphenoid canal area in ventral medial view; (E^1^) explanatory drawing. Not to scale. *Key*: **1**, exit of the MBA(?) through the foramen ovale; **2**, exit of the mPA through the common opening of the alisphenoid canal; **a**, dotted circle marks the same position of the foramen and fenestra in the different aspects; **alc M3**, alveolar capsule of M3; **asc-c**, caudal opening of the alisphenoid canal; **asc-r**, rostral opening of the alisphenoid canal; **bs**, basisphenoid; **bsbp**, basisphenoidal process of the palatine; **fo**, foramen ovale; **IFA**, infraorbital branch of the maxillary artery; **mapf**, major palatine foramen; **MBA(?)**, mandibular branch of the maxillary artery (requires special study); **MPA**, major palatine branch of the maxillary artery (index R means the Avizo segmentation); **mPA**, minor palatine branch of the maxillary artery; **pc**, pterygoid canal; **pf**, piriform fenestra; **pplf**, pterygopalatine fossa; **ps**, presphenoid; **sof-m**, sphenorbital fissure, lateral margin; **tmb**, tympanic bulla.

## Appendix 7

Position of the pterygopalatine fossa in Cricetinae and Arvicolinae. The skull is shown in lateral view with the left part digitally removed. Homology of pplf determined according to Evans (1993: 159). (A) Skull of *Cricetulus barabensis* (ZIN 101697) in dorsal view; (B) the skull of *C. barabensis* in lateral view (cropped); (C^1^) enlarged part of the hamster skull, in lateral view;

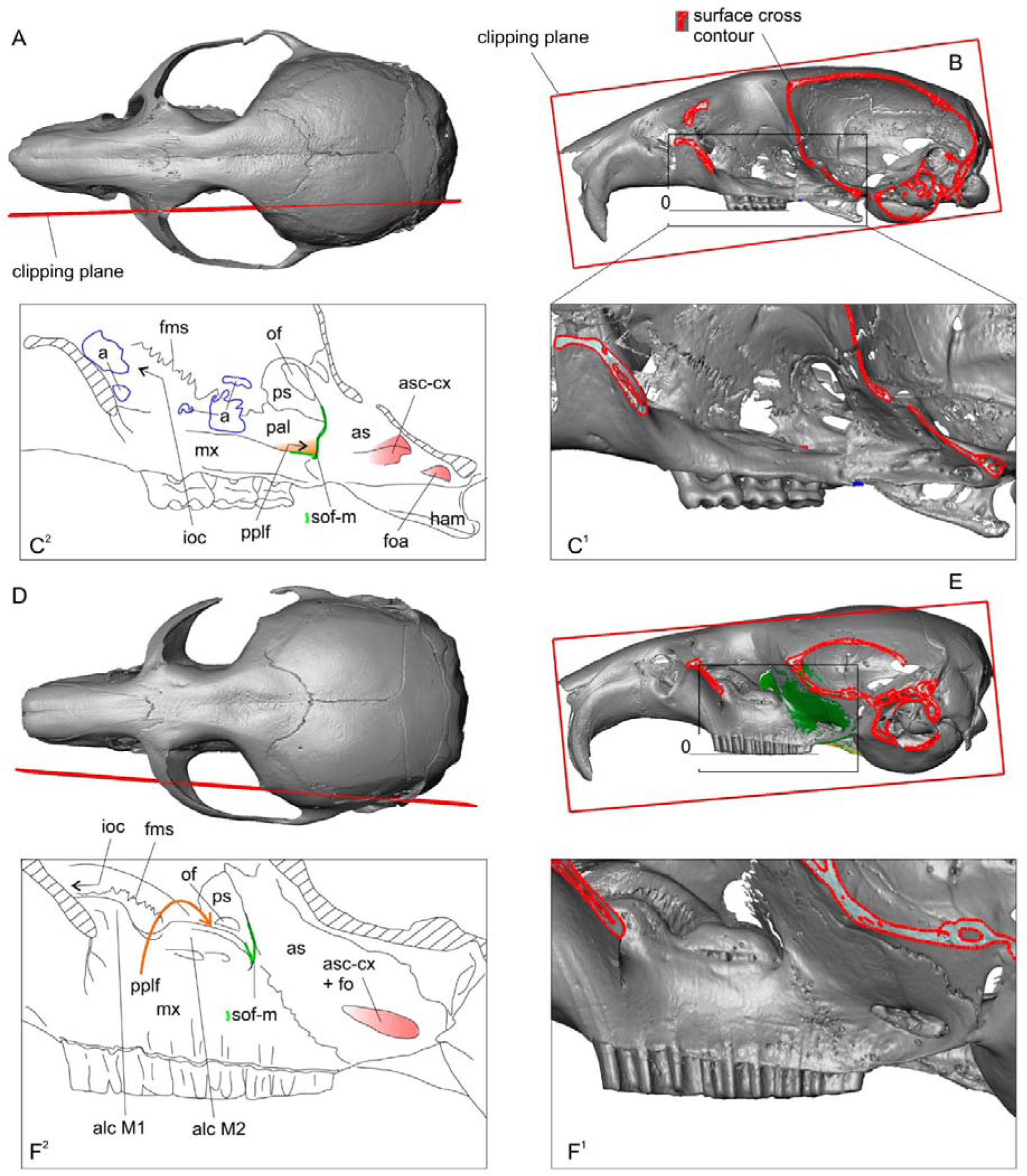

(C^2^) explanatory drawing of the C^1^ picture; (D) skull of *Alticola argentatus* (ZIN 77489) in dorsal view; (E) the skull of *A. argentatus* in lateral view (cropped); (F^1^) enlarged part of the vole skull, in lateral view; (F^2^) explanatory drawing. Not scaled. *Key*: **a**, micro-CT artefacts; **alc**, alveolar capsule of M1 and M2; **as**, alisphenoid; **asc-cx**, alisphenoid canal, complex; **fms**, frontomaxillary suture; **fo**, foramen ovale; **foa**, foramen ovale accessorius; **ham**, pterygoid hamulus; **iof**, infraorbital canal; **mx**, maxilla; **of**, optic foramen; **pal**, palatine; **pplf**, pterygopalatine fossa (orange in C^2^; orange arrow marks a pplf behind alc M2 in F^2^); **ps**, presphenoid; **sof-m**, sphenorbital fissure, margin (light green). Hatched areas indicate the clipping plane.

## Appendix 8

Reconstruction of the passage of the major and minor palatine branches of the internal maxillary artery (detailed Avizo segmentation) of *Cricetulus barabensis* (ZIN 101697) and homology of the foramina of the alisphenoid canal complex of *C. barabensis* and *Neotoma mexicana* (ZIN 39375).

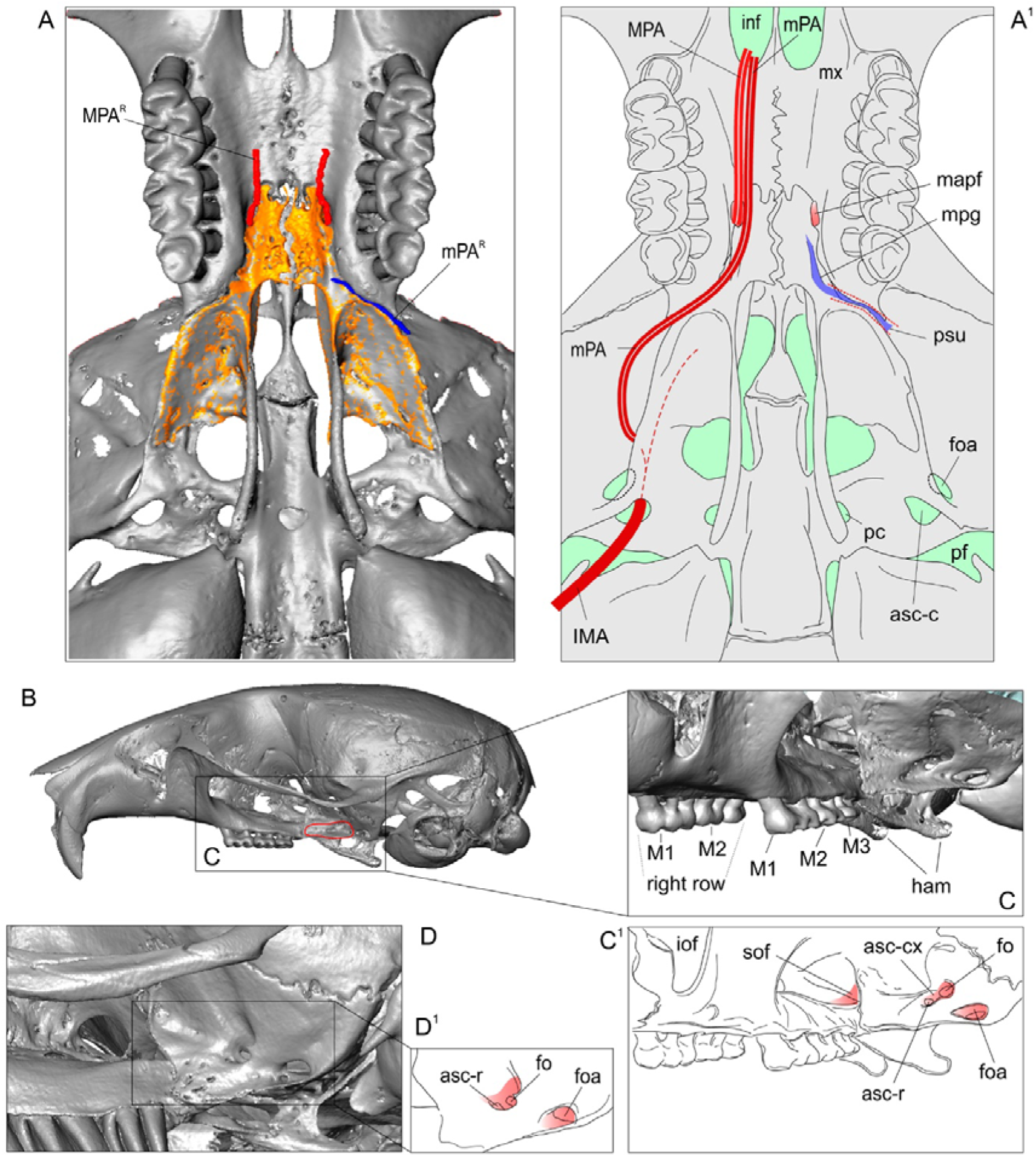

(A) Enlarged part of the skull of *C. barabensis* in ventral view with reconstructed position of the major palatine artery (red) and the minor palatine artery at the entrance to the palatal space; (A1) explanatory drawing; (B) lateral view of the skull of *C. barabensis* (the facet of the external pterygoid muscle is marked in red); (C) enlarged part of the skull of *C. barabensis* in lateral anterior view; (C1) explanatory drawing; (D) enlarged part of the skull of *N. mexicana* in lateral anterior view; (D1) explanatory drawing. Not to scale. *Key*: **asc-c**, caudal opening of the alisphenoid canal; **asc-cx**, alisphenoid canal complex; **asc-r**, rostral opening of the alisphenoid canal; **foa**, foramen ovale accessorius; **ham**, pterygoid hamulus; **IMA**, internal maxillary artery; **inf**, incisive foramen; **iof**, infraorbital foramen; **mapf**, major palatine foramen; **MPA**, major palatine branch of the maxillary artery (index R means the Avizo segmentation); **mPA**, minor palatine branch of the maxillary artery; **mpg**, minor palatine groove; **mx**, maxilla; **pc**, pterygoid canal (posterior opening); **pf**, piriform fenestra; **psu**, groove/canal of the minor palatine artery.

## Appendix 9

The scatter plots show the results of the PSP complex shape analysis with six ’trait vectors’ marked - six groups of paired shape descriptions (I–VI). The reference species are highlighted by the coloured bars and marked by the stars in the legend (C). (A) Morphospace along PC1 and PC2; (B) morphospace along PC3 and PC4; (C) legend. *Key*: **a**, shape transformation trajectory within *Mynomes* species along PC3 from ’narrow-headed’ *M. miurus* to other species of the genus; **b**, similar trajectory within *Lasiopodomys* species along PC3 from true narrow-headed *L. gregalis*/*L. raddei* to *L. brandtii* and *L. mandarinus*.

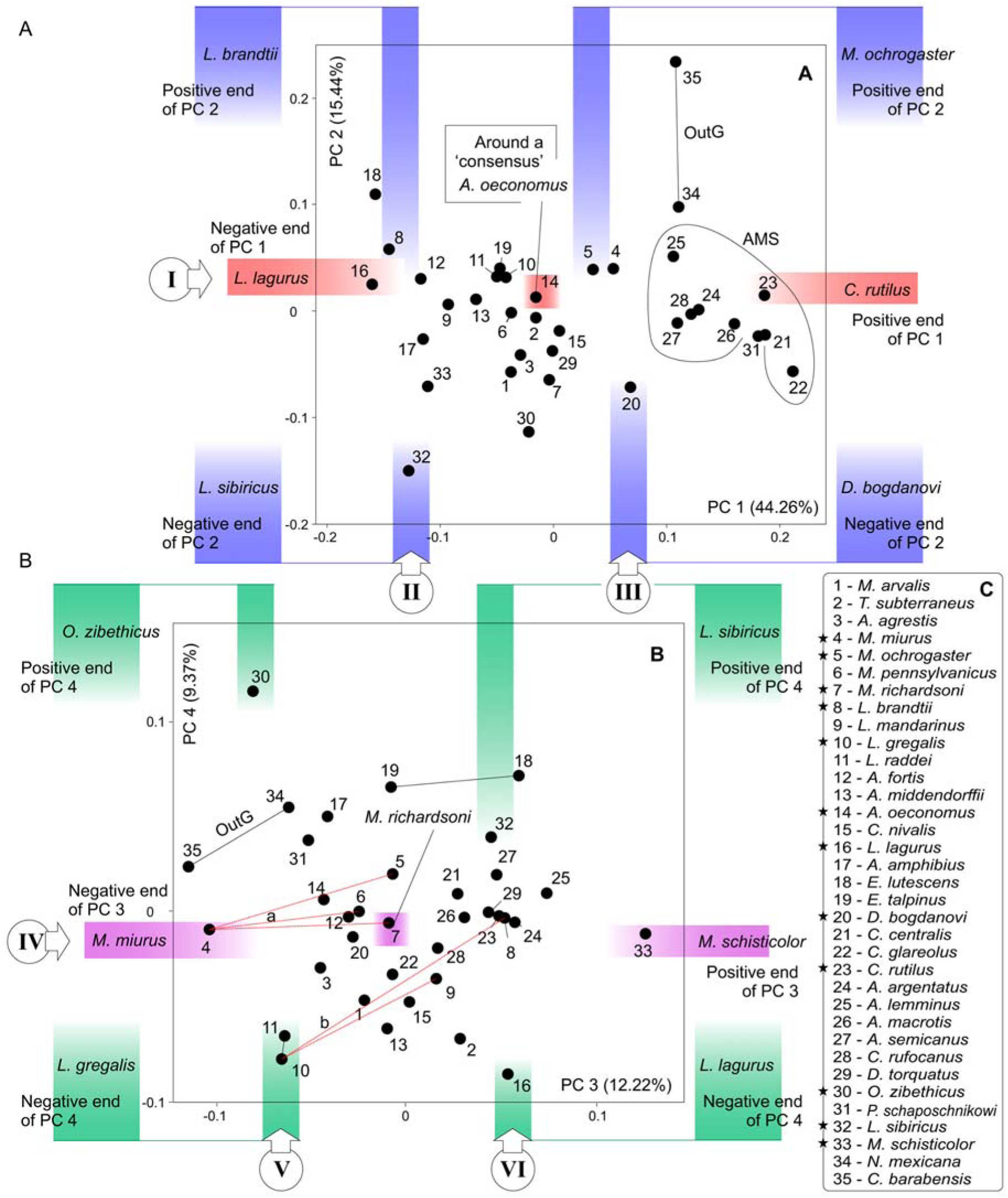

## Appendix 10

Results of the pairwise comparison of 3D models of the PSP complex according to the description of the shape transformations along the III, V and VI trait vectors (see Appendix 9). See also figure 8 in the main text. (A^1^) Pairwise comparison of *M. ochrogaster* and *D. bogdanovi* (trait vector III); the PSP complex shown in ventral posterior aspect; (A^2^) same comparison, the complex shown in dorsal aspect (from the surface of the cranial cavity); (A^3^) plate with statistics: total variance of PC2 and variance absorbed by the third vector (see Appendix 11);

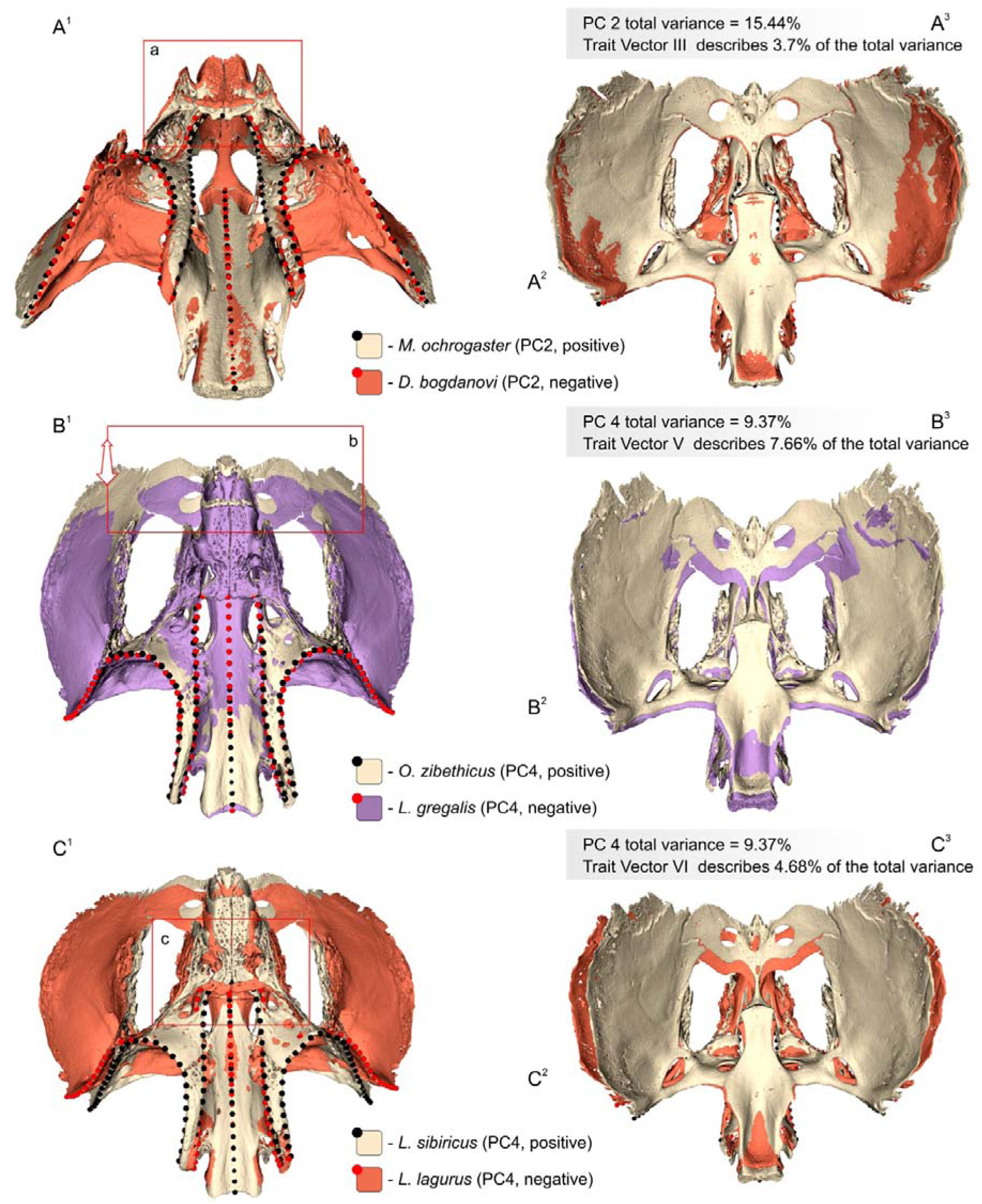

(B^1^) Pairwise comparison of *O. zibethicus* and *L. gregalis* (trait vector V); the PSP complex shown in ventral aspect; (B^2^) same comparison, the complex shown in dorsal aspect; (B^3^) plate with statistics: total variance of PC4 and variance absorbed by the fifth vector; (C^1^) Pairwise comparison of *L. sibiricus* and *L. lagurus* (trait vector VI); the PSP complex shown in ventral aspect; (C^2^) same comparison, the complex shown in dorsal aspect; (C^3^) plate with statistics: total variance of PC4 and variance absorbed by the sixth vector. Not to scale. *Key*: **a**, trait vector III [vIII] — disparity in the horizontal lamina length; **b**, v-V — the transformation in the relative position of the orbital wing of the presphenoid with the relatively stable position of the presphenoid body and the rostral part of the palatine (see Figure 8: C-g) against the background of the ‘narrow’ cranial cavity in the negative area of the PC3 (Figure 8C); **c**, v-VI — disparity in the horizontal lamina length and width.

## Appendix 11

Graphical explanation of the approach to estimating the variance of trait vectors. The approach considers the ratio of the respective pairwise comparison variance (e.g., pairwise comparison of the shapes of *Lagurus* and *Clethrionomys*) to the total variance of each axis (e.g., PC1). The convex hulls at the bottom of the figure are a schematic drawing of the ’space width’ and are taken from figure 7A (see main text). The example calculation was obtained using table 1 and appendix 2.

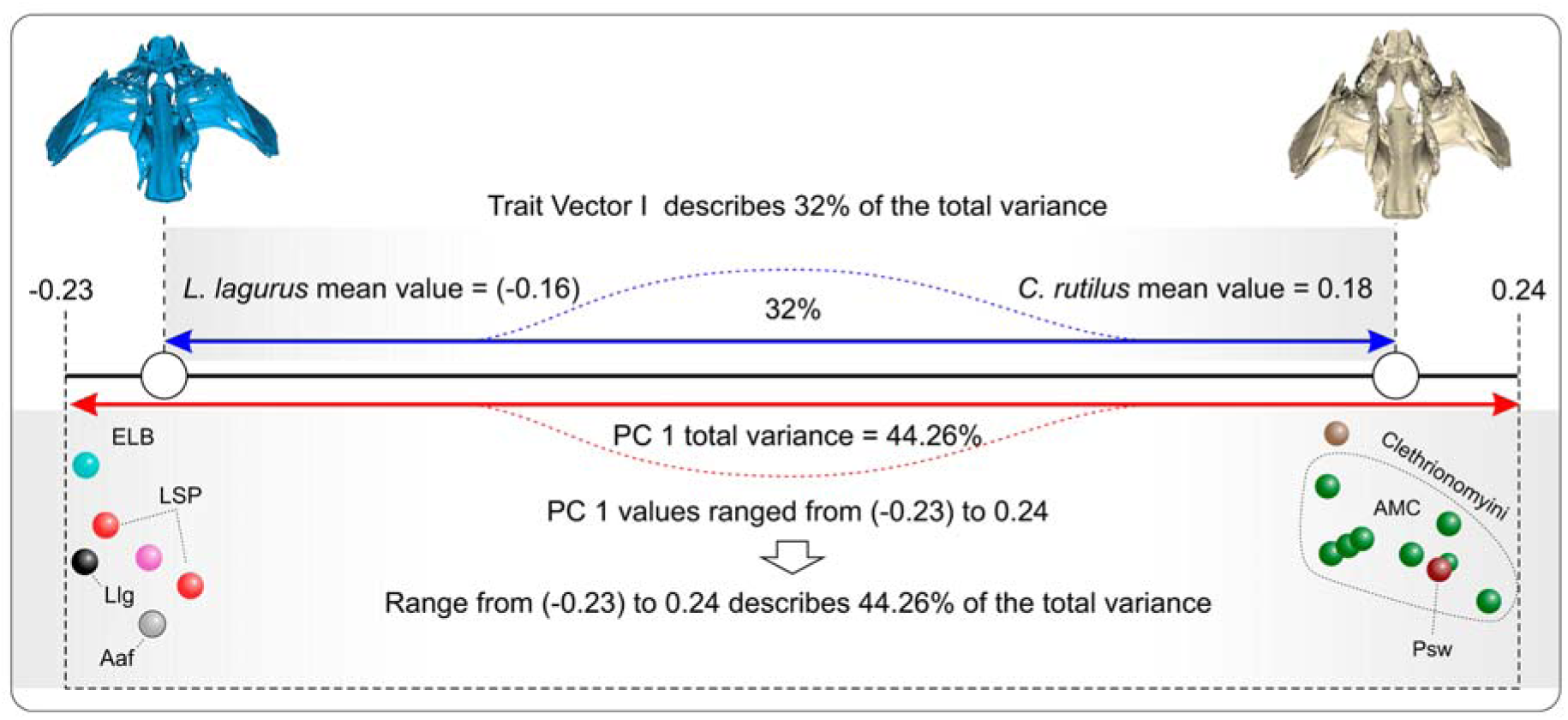

## References

Abramson NI, Bodrov SYu, Bondareva OV, Genelt-Yanovskiy EA, Petrova TV. 2021. A mitochondrial genome phylogeny of voles and lemmings (Rodentia: Arvicolinae): Evolutionary and taxonomic implications. PLoS ONE. 16:e0248198.

Abramson NI, Golenishchev FN, Bodrov SYu, Bondareva OV, Genelt-Yanovskiy EA, Petrova TV. 2020. Phylogenetic relationships and taxonomic position of genus *Hyperacrius* (Rodentia: Arvicolinae) from Kashmir based on evidences from analysis of mitochondrial genome and study of skull morphology. PeerJ. 8:e10364.

Abramson NI, Lebedev VS, Tesakov AS, Bannikova AA. 2009. Supraspecies relationships in the subfamilyArvicolinae (Rodentia, Cricetidae): An unexpected result of nuclear gene analysis. Mol. Biol. 43:834–846.

Adams D.C. 2014. A generalized K statistic for estimating phylogenetic signal from shape and other high-dimensional multivariate data. Syst. Biol. 63:685–697.

Adler D, Murdoch D. 2023. Package ‘rgl’, Version 1.2.8; CRAN, 2023. Available online: https://cran.r-project.org/web/packages/rgl/rgl.pdf (accessed on 07 August 2024)

Bannikova AA, Lebedev VS, Lissovsky AA, Matrosova V, Abramson NI, Obolenskaya EV, Tesakov AS. 2010. Molecular phylogeny and evolution of the Asian lineage of vole genus *Microtus* (Rodentia: Arvicolinae) inferred from mitochondrial cytochrome b sequence. Biol. J. Linn. Soc. 99:595–613.

Baverstock H, Jeffery NS, Cobb SN. 2013. The morphology of the mouse masticatory musculature. J. Anat. 223:46–60.

Borges R, Machado JP, Gomes C, Rocha AP, Antunes A. 2019. Measuring phylogenetic signal between categorical traits and phylogenies. Bioinformatics. 35(11):1862–1869.

Brassard C, Merlin M, Guintard C, Monchâtre-Leroy E, Barrat J, Callou C, Cornette R, Herrel A. 2020a. Interrelations between the cranium, the mandible and muscle architecture in modern domestic dogs. Evol. Biol. 47:308–324.

Brassard C, Merlin M, Guintard C, Monchâtre-Leroy E, Guintard C, Barrat J, Callou C, Cornette R, Herrel A. 2020b. How does masticatory muscle architecture covary with mandibular shape in domestic dogs? Evol. Biol. 47:1.

Brylski P. 1990. Development and evolution of the carotid circulation in geomyoid rodents in relationship to their craniomorphology. J. Morphol. 204:33–45.

Bugge J. 1970. The contribution of the stapedial artery to the cephalic arterial supply in muroid rodents. Acta Anat. 76:313–336.

Christmas MJ, Kaplow IM, Genereux DP, Dong MX, Hughes MX, et al. 2023. Evolutionary constraint and innovation across hundreds of placental mammals. Science. 380:6643.

Claude J. 2008. Morphometrics with R. New York: Springer New York.

Díez del Molino D, Dehasque M, Chacón-Duque JC, Pečnerová P, Tikhonov A, Protopopov A, Plotnikov V, Kanellidou F, Nikolskiy P, Mortensen P, Danilov GK, Vartanyan S, Gilbert MTP, Lister AM, Heintzman PD, van der Valk T, Dalén L. 2023. Genomics of adaptive evolution in the woolly mammoth. Curr. Biol. 33:1–12.

Drake AG. 2011. Dispelling dog dogma: an investigation of heterochrony in dogs using 3D geometric morphometric analysis of skull shape. Evol. Dev. 13:204–213.

Evans HE. 1993. Miller’s anatomy of the dog. Philadelphia: W.B. Saunders.

Fedorov A, Beichel R, Kalpathy-Cramer J, Finet J, Fillion-Robin J-C, Pujol S, Bauer C, Jennings D, Fennessy FM, Sonka M, et al. 2012. 3D Slicer as an image computing platform for the quantitative imaging network. Magn. Reson. Imaging. 30:1323–1341.

Fejfar O, Heinrich W-D, Kordos L, Maul LC. 2011. Microtoid cricetids and the early hystory of arvicolids (Mammalia, Rodentia). Palaeontol. Electron. 14.3.27A.

Fabre P-H, Cornette R, Perrard A, Boyer DM, Prasad GVR, Hooker JJ, Goswami A. 2014. A Three-dimensional morphometric analysis of the locomotory ecology of *Deccanolestes*, a eutherian mammal from the Late Cretaceous of India. J. Vertebr. Paleontol. 34:146–156.

Fabre P-H, Reeve AH, Fitriana YS, Aplin KP, Helgen KM. 2018. A new species of *Halmaheramys* (Rodentia: Muridae) from Bisa and Obi Islands (North Maluku Province, Indonesia). J. Mammal. 99:187–208.

Ginot S, Claude J, Hautier L. 2018a. One skull to rule them all? Descriptive and comparative anatomy of the masticatory apparatus in five mouse species. J. Morphol. 279(9): 1234– 1255.

Ginot S, Herrel A, Claude J, Hautier L. 2018b. Skull size and biomechanics are good estimators of in vivo bite force in murid rodents. Anat. Rec. 301:256–266.

Gromov IM, Erbaeva MA. 1995.Mammals of the fauna of Russia and the adjacent territories. Lagomorphs and Rodents (the Identification Guide, Issue 167). Saint-Petersburg: Zoological Institute of the Russian academy of sciences. [in Russian]

Hammer Ø, Harper DAT, Ryan PD. 2001. PAST: Paleontological statistics soft-ware package for and data analysis. Palaeontol. Electron. 4:1–9.

Hautier L. 2010. Masticatory muscle architecture in the gundi *Ctenodactylus vali* (Mammalia, Rodentia). Mammalia. 74:153–162.

Hiller M, Schaar BT, Indjeian VB, Kingsley DM, Hagey LR, Bejerano G. 2012. A “forward genomics” approach links genotype to phenotype using independent phenotypic losses among related species. Cell Rep. 2:817–823.

Horn JL. 1965. A rationale and test for the number of factors in factor analysis. Psychometrica. 30:179–185.

Hu Z, Sackton TB, Edwards SV, Liu JS. 2019. Bayesian detection of convergent rate changes of conserved noncoding elements on phylogenetic trees. Mol. Biol. Evol. 36:1086–1100.

Jackson DA. 1993. Stopping rules in principal components analysis: A comparison of heuristical and statistical approaches. Ecology. 74:2204–2214.

Kamilar JM, Cooper N. 2013. Phylogenetic signal in primate behaviour, ecology and life history. Philos. Trans. R. Soc. B Biol. Sci. 368: 20120341–20120341.

Kesner MH. 1980. Functional morphology of the masticatory musculature of the rodent subfamily Microtinae. J. Morphol. 165:205–222.

Klingenberg CP. 2009. Morphometric integration and modularity in configurations of landmarks: tools for evaluating a priori hypotheses. Evol. Dev. 11(4):405–421.

Koenigswald von W. 1993. Heterochronies in morphology and schmelzmuster of hypsodont molars in the Muroidea (Rodentia). Quat. Int. 19:57–61.

Kowalczyk A, Meyer WK, Partha R, Mao W, Clark NL, Chikina M. 2019. RERconverge: an R package for associating evolutionary rates with convergent traits. Bioinformatics. 35:4815– 4817.

Krause DW, Hoffmann S, Wible JR, Kirk EC, Schultz JA, von Koenigswald W, Groenke JR, Rossie JB, O’Connor PM, Seiffert ER, Dumont ER, Holloway WL, Rogers RR, Rahantarisoa LJ, Kemp AD, Andriamialison H. 2014. First cranial remains of a gondwanatherian mammal reveal remarkable mosaicism. Nature. 515:512–517.

Kryštufek B, Shenbrot GI. 2022. Voes and lemmings (Arvicolinae) of the Palaearctic Region. Maribor: University of Maribor Press.

Lartillot N, Poujol R. 2011. A phylogenetic model for investigating correlated evolution of substitution rates and continuous phenotypic characters. Mol. Biol. Evol. 28:729–744.

Ligges U, Maechler M, Schnackenberg S. 2023. Package ‘scatterplot3d’, Version 0.3-44; CRAN, 2023. Available online: https://cran.r-project.org/web/packages/scatterplot3d/scatterplot3d.pdf (accessed on 07 August 2024)

MacPhee RDE. 1981. Auditory regions of primates and eutherian insectivores: morphology, ontogeny and character analysis. Contrib. Primatol. 18:1–282.

MacPhee RDE. 1994. Morphology, adaptations, and relationships of *Plesiorycteropus*, and a diagnosis of a new order of eutherian mammals. Bull. Am. Mus. Nat. Hist. 220:1–214.

Mao F, Zhang C, Ren J, Wang T, Wang G, Zhang F, Rich T, Vickers-Rich P, Meng J. 2024. Fossils document evolutionary changes of jaw joint to mammalian middle ear. Nature. 628:576–581.

Maul LC. 2001. The transition from hypsodonty to hypselodonty in the *Mimomys savini* – *Arvicola* lineage. Lynx. 32: 247–253.

McDowell SB Jr, 1958. The Greater Antillean insectivores. Bull. Am. Mus. Nat. Hist. 115(3):113–214.

Missagia RV, Perini FA. 2018. Skull morphology of the Brazilian shrew mouse *Blarinomys breviceps* (Akodontini; Sigmodontinae), with comparative notes on Akodontini rodents. Zool. Anz. 277:148–161.

Michaux J, Hautier L, Hutterer R, Lebrun R, Guy F, García-Talavera F. 2012. Body shape and life style of the extinct rodent *Canariomys bravoi* (Mammalia, Murinae) from Tenerife, Canary Islands (Spain). C. R. Palevol. 11:485–494.

Monroy PLC, Grefte S, Kuijpers-Jagtman AM, Helmich MPAC, Ulrich DJO, Von den Hoff JW, Wagener FADTG. 2013. A rat model for muscle regeneration in the soft palate. PLoS ONE 8:e59193.

Musser GG, Carleton MD. 2005. Superfamily Muroidea. In: Wilson DE, Reeder DA, editors. Mammal species of the world: a taxonomical reference. 3rd edition. Vol. 2. Baltimore: Johns Hopkins University Press; p. 894–1531.

Novacek MJ. 1986. The skull of leptictid insectivorans and the higher-level classification of eutherian mammals. Bull. Am. Mus. Nat. Hist. 183(1):1–112.

Pardiñas UFJ, Myers P, León-Paniagua L, Ordґóñez Garza N, Cook JA, Kryštufek B, Cook J, Salazar-Bravo J. 2017. Family Cricetidae (true hamsters, voles, lemmings and new world rats and mice). In: Wilson DE, Lacher Jr. TE, Mittermeier RA, editors. Handbook of the mammals of the world. Vol. 7. Barcelona: Lynx Edicions; p. 204–279.

Petrova T, Skazina M, Kuksin A, Bondareva O, Abramson N. 2022. Narrow-headed voles species complex (Cricetidae, Rodentia): Evidence for species differentiation inferred from transcriptome data. Diversity. 14:512.

Pozdnyakov AA. 2008. The bony palate morphology in Arvicolinae (Rodentia: Cricetidae), with comments on taxonomy and nomenclature. Sbornik trudov Zoologicheskogo muzeya MGU 49:184–209, [in Russian, with English summary].

Prudent X, Parra G, Schwede P, Roscito JG, Hiller M. 2016. Controlling for phylogenetic relatedness and evolutionary rates improves the discovery of associations between species’ phenotypic and genomic differences. Mol. Biol. Evol. 33: 2135–2150.

Quay WB. 1954. The anatomy of the diastemal palate in microtine rodents. Misc. Pub. Mus. Zool. U. Michigan. 86:1–49.

Redlich R, Kowalczyk A, Tene M, Sestili HH, Foley K, Saputra E, Clark N, Chikina M, Meyer WK, Pfenning A. 2023. RERconverge expansion: using relative evolutionary rates to study complex categorical trait evolution. bioRxiv. 10.1101/2023.12.06.570425

Renaud S, Alibert P, Auffray J-C. 2012. Modularity as a source of new morphological variation in the mandible of hybrid mice. BMC Evol. Biol. 12:141.

Repenning CA. 1968. Mandibular musculature and the orifgin of the subfamily Arvicolinae (Rodentia). Acta Zool. Crac. 13:29–72.

Robovský J, ŘIčánková V, Zrzavý J. 2008. Phylogeny of Arvicolinae (Mammalia, Cricetidae): utility of morphological and molecular data sets in a recently radiating clade. Zool. Scr. 37:571–590.

Rohlf FJ. 1993. Relative warp analysis and an example of its application to mosquito wings. In: Marcus LF, Bello E, Garcia-Valdecasas A, editors. Contributions to morphometrics. Monografias 8. Madrid: Museo Nacional de Ciencias Naturales; p. 131–159.

Rohlf FJ, Slice DE. 1990. Extensions of the Procrustes method for the optimal superimposition of landmarks. Syst. Zool. 39:40–59.

Schlager S. 2017. Morpho and Rvcg – shape analysis in R: R-packages for geometric morphometrics, shape analysis and surface manipulations. In: Zheng G, Li S, Szekely G, editors. Statistical Shape and Deformation Analysis, 1st. Edition. San Diego: Academic Press Inc.; p. 217–256.

Singh N, Albert FW, Plyusnina I, Trut L, Pääbo S, Harvati K. 2017. Facial shape differences between rats selected for tame and aggressive behaviors. PLoS ONE. 12:e0175043.

Smith AB. 1994. Systematics and the fossil record: documenting evolutionary patterns. Oxford: Blackwell Science Ltd.

Terray L, Denys C, Goodman SM, Soarimalala V, Lalis A, Cornette R. 2022. Skull morphological evolution in Malagasy endemic Nesomyinae rodents. PLoS ONE. 17:e0263045.

Van Valen LM. 1994. Serial homology: the crests and cusps of mammalian teeth. Acta Palaeontol. Pol. 38:145–158.

Voet I., Denys C, Colyn M, Lalis A, Konečny A, Dlapré A, Nicolas V, Cornette, R. 2022. Incongruences between morphology and molecular phylogeny provide an insight into the diversifcation of the *Crocidura poensis* species complex. Sci. Rep. 2:e10531.

Vorontsov NN. 1982. The Lower hamster (Cricetidae) of the world fauna. Part 1: Morphology and ecology. Leningrad: Nauka. [In Russian]

Voyta LL, Abramov AV, Lavrenchenko LA, Nicolas V, Petrova EA, Kryuchkova LYu. 2022a. Dental polymorphisms in *Crocidura* (Soricomorpha: Soricidae) and evolutionary diversification of crocidurine shrew dentition. Zool. J. Linn. Soc. 196:1069–1093.

Voyta LL, Izvarin EP, Shemyakina YA, Nikiforova VA, Strukova TV, Smirnov NG, Melnikov DA, Bobretsov AV. 2023. Morphospace dynamics and interspecies variety of *Sorex araneus* and *S. tundrensis* according to recent and fossil data. Palaeontol. Electron. 26(3):a51.

Voyta LL, Omelko VE, Tiunov MP, Petrova EA, Kryuchkova LYu. 2022b. Temporal variation in soricid dentition: which are first – qualitative or quantitative features? Hist. Biol. 34(10):1901–1915.

Voyta LL, Omelko VE, Tiunov MP, Vinokurova MA. 2021. When beremendiin shrews disappeared in East Asia, or how we can estimate fossil redeposition. Hist. Biol. 33(11):2656–2667.

Voyta LL, Petrova TV, Panitsina VA, Bodrov SYu, Winkler V, Kryuchkova LYu, Bramson NI. 2024. A Cybertaxonomic revision of the “Crocidura pergrisea” species complex with a special focus on endemic rocky shrews: *Crocidura armenica* and *Crocidura arispa* (Soricidae). Biology. 13(6):448.

Wang H, Meng J, Wang Y. 2019. Cretaceous fossil reveals a new pattern in mammalian middle ear evolution. Nature. 576:102–105.

Weksler M. 2006. Phylogenetic relationship of oryzomine rodents (Muroidea: Sigmodontinae): separate and combined analyses of morphological and molecular data. Bull. Am. Mus. Nat. Hist. 296(149):1–48.

Wible JR. 2008. On the cranial osteology of the Hyspaniolan solenodon, *Solenodon paradoxus* Brandt, 1833 (Mammalia, Lipotyphla, Solenodontidae). Ann. Carnegie Mus. 77:321–402.

Wible JR. 2022. CT study of the cranial osteology of the Gray short-tailed opossum *Monodelphisdomestica* (Wagner, 1842) (Marsupialia, Didelphidae) and comments on the internal nasal skeleton floor. Ann. Carnegie Mus. 87:249–289.

Wible JR, Wang Y, Li C, Dawson MR. 2005. Cranial anatomy and relationships of a new ctenodactyloid (Mammalia, Rodentia) from the Early Eocene of Hubei Province, China. Ann. Carnegie Mus. 74:91–150.

Withnell CB, Scarpetta SG. 2024. A new perspective on the taxonomy and systematics of Arvicolinae (Gray, 1821) and a new time-calibrated phylogeny for the clade. PeerJ. 12:e16693.

Young RL, Badyaev AV. 2010. Developmental plasticity links local adaptation and evolutionary diversification in foraging morphology. J. Exp. Zool. B: Mol. Dev. Evol. 314:434–444.

Zazhigin VS. 1980. Late Pliocene and Anthropogene rodents of the South of Western Siberia. Moscow: Nauka. [In Russian].

Zelditch ML, Swiderski DL, Sheets HD, Fink WL. 2004. Geometric morphometrics for biologists: A Primer. Amsterdam: Elsevier Academic Press.

